# Precise modulation of transcription factor levels reveals drivers of dosage sensitivity

**DOI:** 10.1101/2022.06.13.495964

**Authors:** Sahin Naqvi, Seungsoo Kim, Hanne Hoskens, Harry S. Matthews, Richard A. Spritz, Ophir D. Klein, Benedikt Hallgrímsson, Tomek Swigut, Peter Claes, Jonathan K. Pritchard, Joanna Wysocka

## Abstract

Transcriptional regulation displays extensive robustness, but human genetics indicate sensitivity to transcription factor (TF) dosage. Reconciling such observations requires quantitative studies of TF dosage effects at trait-relevant ranges, which are lacking to date. TFs play central roles in both normal-range and disease-associated variation in facial morphology; we therefore developed an approach to precisely modulate TF levels in human facial progenitors and applied it to SOX9, a TF associated with craniofacial variation and disease (Pierre Robin Sequence, PRS). We found that most SOX9-dependent regulatory elements (REs) are buffered against small decreases in SOX9 dosage, but REs directly and primarily regulated by SOX9 show heightened sensitivity to SOX9 dosage; these RE responses predict gene expression responses. Sensitive REs and genes underlie the vulnerability of chondrogenesis and PRS-like craniofacial shape variation to SOX9 dosage perturbation. We propose that such REs and genes drive the sensitivity of specific phenotypes to TF dosage, while buffering of other genes leads to robust, nonlinear dosage-to-phenotype relationships.

## INTRODUCTION

Transcriptional regulation is fundamental to the control of gene expression, and is mediated by sequence-specific transcription factors (TFs), a class of proteins that modulate target gene expression by binding to specific DNA motifs within noncoding regulatory elements (REs); TFs are thus the major drivers of cellular and developmental identity^1^. As evidenced by the stability of organismal development despite environmental and genetic variation^2^, cellular and developmental programs must be robust to fluctuations in TF levels. Additional evidence indicates that cis-regulatory landscapes are also robust to perturbations, with loss of individual REs often leading to minimal effects on gene expression and/or morphological phenotypes^3, 4^. Many naturally-occurring genetic variants in TF motifs also do not result in changes in TF binding or gene expression^5, 6^.

Despite such robustness, human genetic studies indicate phenotypic sensitivity to TF dosage. For instance, TFs are strongly enriched for haploinsufficient disease associations, resulting from the loss of one functional allele and ∼50% dosage reduction, and are depleted of loss-of-function (LoF) variants in the general population ^7, 8^. Genome-wide association studies have revealed thousands of trait-associated common variants, many of which likely act by modulating RE activity and target gene expression levels^9, 10^; trait-associated variants are also highly enriched around TF genes^11, 12^. Both experimental and population-level data generally suggest per-allele effects of up to 10-15%^13, 14^, which may be even smaller for important, dosage-sensitive genes such as TFs. Together, available evidence indicates that small, <50% variation in TF levels leads to variation in complex traits or common disease risk, while ∼50% reductions in TF dosage can lead to severe disorders.

Understanding how cellular and developmental programs can be simultaneously robust and sensitive to TF levels is a fundamental problem, and requires systematic, quantitative studies of the effects of endogenous TF dosage changes at physiologically relevant levels. However, most studies of TF function to date have used knockouts, overexpression resulting in dosage increases far beyond the ranges relevant for trait variation and disease, or genome-wide assays of TF binding in unperturbed states. Such studies have found that TFs typically regulate hundreds to thousands of REs and genes^15–18^, and when knocked out in a developmental context, produce pleiotropic, often embryonic lethal, phenotypes. Nonlinearity in the effects of TF dosage have been proposed to underlie TF haploinsufficiency^19, 20^, an idea based on Fisher’s 1931 model for dominance^21^, but such models have not been directly tested experimentally.

The importance of transcriptional regulation is apparent in the development of the human face, which forms a key component of individual physical identity and inter-personal interactions and is disrupted in a broad range of craniofacial disorders that together account for approximately one-third of birth defects^22^. Much of the normal range of facial shape as well as disease-associated variation derives from cranial neural crest cells (CNCCs), a transient, embryonic cell population that arises from the neural folds, and migrates ventrally to the developing facial prominences, later giving rise to most of the craniofacial skeleton and connective tissue^23^. We recently reviewed the genetics of human craniofacial morphology and found that genes encoding TFs are frequently involved in both common (influencing normal-range shape variation) and rare (causative for typically Mendelian and haploinsufficient disorders) variation in craniofacial morphology^24^. Thus, studying the quantitative effects of TF dosage alterations on molecular, cellular, and morphological phenotypes in craniofacial development could provide general insights into mechanisms underlying dosage sensitivity and/or robustness in human phenotypes.

Multiple lines of evidence specifically highlight the developmentally important TF SOX9 as an attractive model for studying TF dosage effects. Heterozygous LoF mutations in *SOX9* cause campomelic dysplasia, which is a disorder manifesting in long-bone defects, male-to-female sex reversal, and a set of craniofacial features termed Pierre Robin Sequence (PRS), characterized by underdevelopment of the lower jaw (micrognathia)^25, 26^. These observations suggest that among the diverse cell types in which SOX9 plays key regulatory roles (reviewed in ^27^), CNCCs, chondrocytes, and Sertoli cells exhibit heightened sensitivity to ∼50% reduction in SOX9 dosage. Isolated PRS without other manifestations of campomelic dysplasia can also be caused by heterozygous deletion of CNCC-specific enhancers of *SOX9* (Benko et al., 2009; Long et al, 2020), whereas common genetic variants in noncoding regions at the *SOX9* locus are associated with normal-range facial variation in humans of both European and East Asian ancestry^29–31^. Furthermore, CNCC-specific modulation of *Sox9* dosage in mice revealed that craniofacial development is sensitive to changes in *Sox9* dosage over a broad range^32^. Even 10-13% reduction in *Sox9* mRNA levels in the developing mouse face is sufficient to produce a subtle but reproducible change in lower jaw morphology^32^.

Here, we sought to understand the response to quantitative changes in SOX9 dosage at multiple levels: chromatin, gene expression, cellular phenotypes, and facial morphology. We applied the degradation tag (dTAG) system to achieve tunable and conditional modulation of SOX9 dosage in an *in vitro* model of human CNCC development. We found the chromatin accessibility response of REs to changes in SOX9 dosage to be broadly buffered against small to moderate changes in SOX9 dosage. However, a subset of REs associated with specific regulatory features shows heightened sensitivity to SOX9 dosage. Gene expression shows a similar, primarily buffered, response to SOX9 dosage, with a subset of sensitive genes; this range of responses can be predicted from changes in chromatin accessibility. Pro-chondrogenic genes, *in vitro* chondrogenesis itself, and genes and REs associated with PRS-like phenotypes exhibit heightened sensitivity to SOX9 dosage. We synthesize these observations into a model in which phenotypically impactful REs and genes sensitive to SOX9 dosage transmit quantitative TF dosage changes to specific cellular and morphological effects underlying craniofacial shape variation and disease, while other phenotypically important REs and genes are regulated by SOX9 but highly buffered.

## RESULTS

### Precise modulation of SOX9 dosage in hESC-derived CNCCs

Based on previous reports that the degradation tag (dTAG) system could be used for rapid or tunable target degradation in human or mouse cells^33–35^, we sought to apply dTAG to modulate SOX9 dosage in human embryonic stem cell (hESC)-derived CNCCs. Briefly, our approach involves genome editing in hESCs to tag the protein of interest (SOX9) with FKBP12^V^^36^, which mediates degradation upon addition of a degrader molecule (dTAG^V^-1), the fluorescent protein mNeonGreen as a quantitative proxy for SOX9 levels, and the V5 epitope tag for biochemical assays. Using a selection-free genome editing method^36^, we obtained two hESC clones with biallelic knock-in of the intact FKBP12^V^^36^-mNeonGreen-V5 tag at the *SOX9* C-terminus (Extended Data Fig. 1A).

To avoid indirect effects of depleting SOX9 during the hESC-to-CNCC differentiation, we first differentiated tagged hESCs into CNCCs using an established protocol with neuroepithelial sphere intermediates^37, 38^ and subsequently titrated SOX9 concentration by adding different dTAG^V^-1 concentrations to CNCCs. We assessed downstream effects on chromatin accessibility and gene expression by the assay for transposase-accessible chromatin (ATAC) and RNA sequencing, respectively (Fig. 1A). Differentiation of *SOX9*-tagged hESCs revealed bright, nuclear fluorescence in a subset of cells within neuroepithelial spheres and continued fluorescence in early-stage migratory CNCC (Fig. 1B), consistent with known roles of SOX9 in neural crest specification and migration^39, 40^. Later-stage *SOX9*-tagged CNCCs showed similar SOX9 levels as untagged (WT) CNCCs (Fig. 1F), and absolute SOX9 levels between the two *SOX9*-tagged clones were very similar when assessed by flow cytometry (Extended Data Fig. 1B). We first treated *SOX9*-tagged CNCCs with a 10-fold dilution series of dTAG^V^-1 for 24h, and observed a gradual change in SOX9 levels (Fig. 1C). Further optimization of dTAG^V^-1 concentrations and extension of treatment time to 48h yielded six distinct and reproducible SOX9 concentrations (Fig. 1D, right). Inspection of the fluorescence distribution in single cells revealed a unimodal distribution that shifted towards lower levels with higher dTAG^V^-1 concentrations, indicating uniform effects of dTAG^V^-1 on SOX9 depletion (Fig. 1D, left). Together, these results indicate that dTAG can be used to precisely modulate SOX9 dosage in hESC-derived CNCCs.

**Figure 1.**
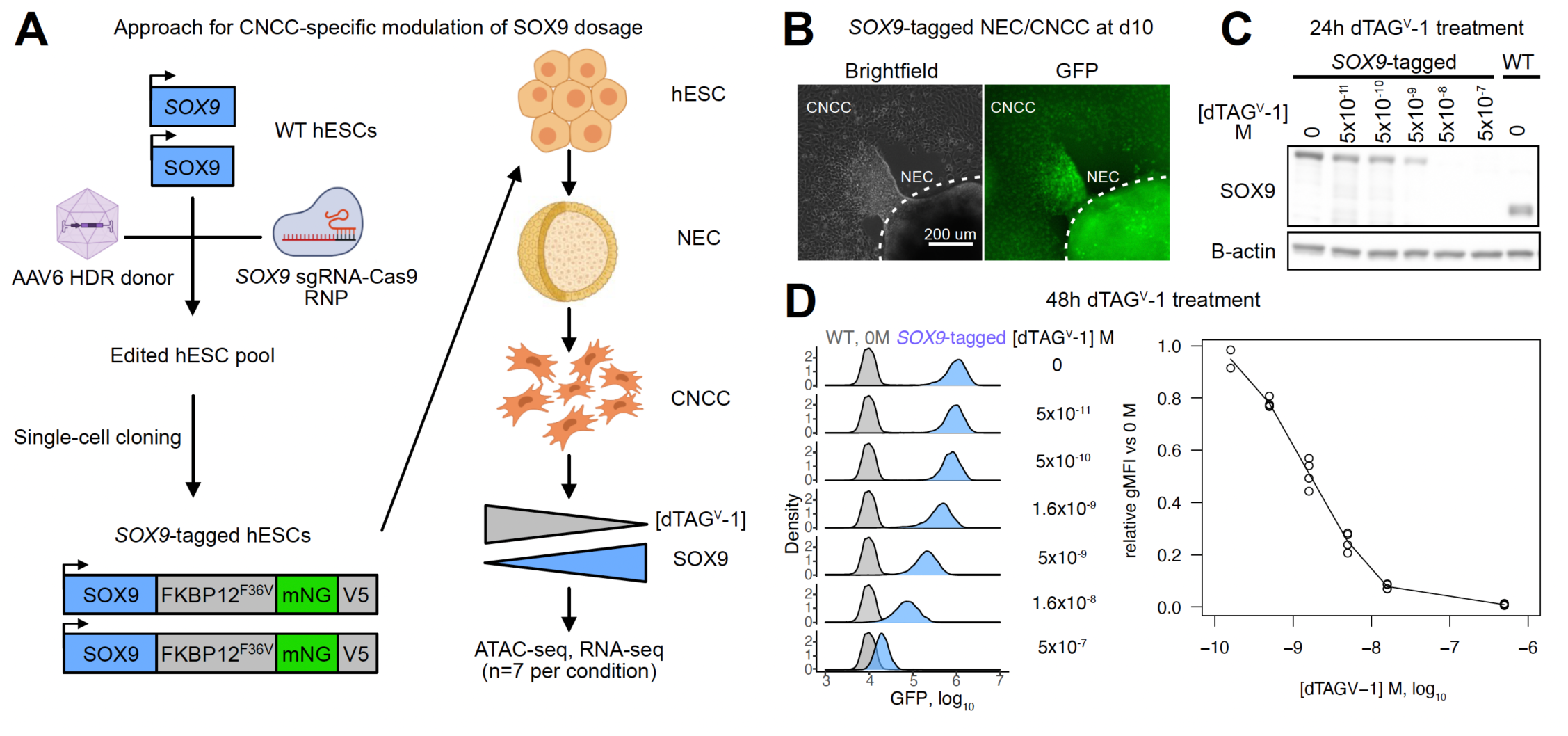
Precise modulation of SOX9 dosage in hESC-derived CNCCs. (A) Schematic of hESC editing and CNCC-specific SOX9 titration approach using dTAG. AAV, adeno-associated virus; HDR, homology-directed repair; RNP, ribonucleoprotein; NEC, neuroectodermal spheres. (B) Live-cell imaging of mNeonGreen fluorescence in attached NEC and migrating CNCC derived from *SOX9*-tagged hESCs at the time of CNCC delamination from neuroepithelial spheres (day 10). (C) Western blot of SOX9 in *SOX9*-tagged or WT passaged mesenchymal CNCCs, treated with indicated concentrations of dTAG^V^-1 over 24h. (D) Flow cytometry analysis of mNeonGreen fluorescence intensity in *SOX9*-tagged CNCCs as a function of increasing dTAG^V^-1 concentrations across single-cells (left) or averaged per biological replicate (differentiation or clone, right). gMFI, geometric mean.

### Most SOX9-dependent REs are buffered in their response to SOX9 dosage changes, with a sensitive subset

To assess the effect of SOX9 dosage changes on chromatin accessibility, we performed ATAC-seq on *SOX9*-tagged CNCCs with six different dTAG^V^-1 concentrations (and therefore at a range of SOX9 levels, Fig. 1D), as well as on WT CNCCs treated with either DMSO or the highest dTAG^V^-1 concentration used (500 nM). Principal component analysis on ATAC counts per million (CPM) values at each of the 151,457 reproducible peak regions (herein referred to as regulatory elements, REs) revealed a clear direction of dTAG^V^-1 (and thus SOX9 titration) effect in principal component space, independent of differentiation batch effects (Extended Data Fig. 2A).

WT CNCCs treated with 500 nM dTAG^V^-1 did not show the same effect in principal component space and had no significantly (5% FDR) changed REs, as compared to 6,169 changed REs from two *SOX9*-tagged replicates treated with 500 nM dTAG^V^-1 (Extended Data Fig. 2B), indicating minimal confounding by off-target effects. We plotted each *SOX9*-tagged sample’s loading in this principal component direction as a function of its estimated SOX9 dosage, revealing a clearly nonlinear relationship, as assessed by a decreased Aikake Information Criterion (AIC, indicating a better fit) for various nonlinear functions as compared to a linear function (Fig. 2A). These results also suggest a largely monotonic effect of SOX9 dosage on individual RE accessibility, which we directly confirmed by pairwise comparisons between all reduced SOX9 dosages and full dosage (Extended Data Fig. 2C,D).

**Figure 2.**
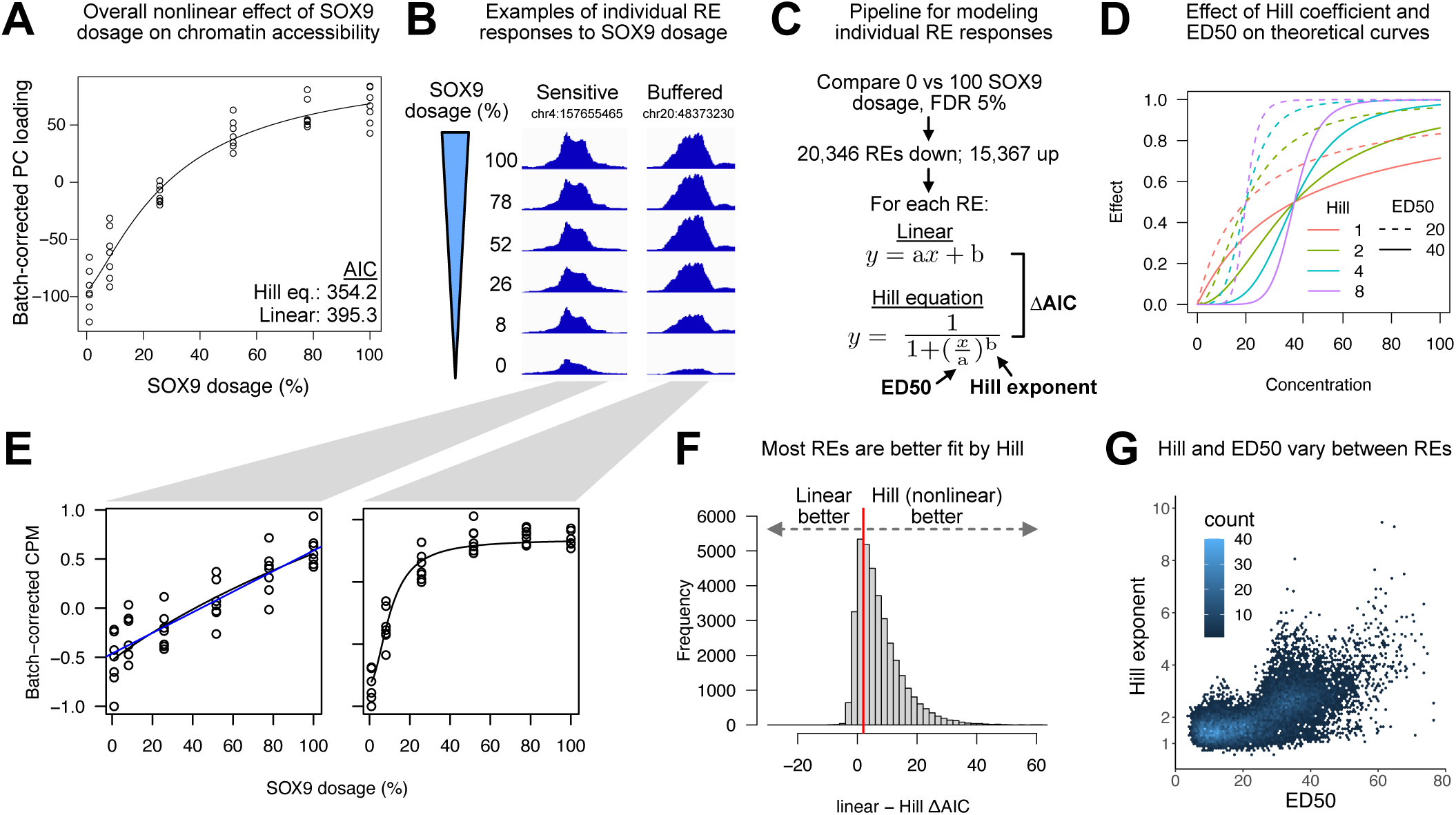
Most SOX9-dependent REs are buffered in their response to SOX9 dosage changes, with a sensitive subset. (A) Loadings from principal component analysis of ATAC-seq counts per million (CPM) of all 151,457 REs across all CNCC samples (see Extended Data Fig. 2A), corrected for differentiation batch and plotted as a function of estimated relative SOX9 dosage (shown as % relative to no dTAG^V^-1). Black line represents Hill equation fit. AIC, Aikake’s Information Criterion. Points are biological (differentiation or clone) replicates. (B) Example ATAC-seq browser tracks of individual RE accessibility at different SOX9 dosages (y-axis), averaged across all replicates at each dosage. (C) Schematic of approach to model nonlinearity of SOX9-dependent REs. (D) Illustration of different ED50 and Hill exponent value on theoretical Hill equation curves. (E) Individual REs from (B) with replicates, fit by linear (blue line) and/or Hill equation (black line). (F) Histogram of τιAIC of all SOX9-dependent REs. Red line indicates τιAIC = 2. (G) Scatterplot of ED50 and Hill exponent across individual SOX9-dependent REs with good fit (p < 0.05 for either parameter).

Inspection of individual REs revealed different responses to SOX9 dosage, with some showing fairly constant decreases in accessibility correlated with SOX9 dosage, and others showing buffered responses (*i.e*., minimal change in accessibility until SOX9 dosage is greatly reduced) (Fig. 2B,E), with a similar variety for REs gaining accessibility upon SOX9 depletion (Extended Data Fig. 2E). To model these responses, we first defined all SOX9-dependent REs as those that respond significantly (5% FDR) to full depletion of SOX9 over 48h, using all seven replicates. This yielded 35,713 SOX9-dependent REs, of which 20,346 decreased and 15,367 increased in accessibility upon SOX9 depletion (Supplementary Table 1). We modeled batch-corrected ATAC CPM values of each SOX9-dependent RE as a function of estimated SOX9 dosage using two functions – 1) linear, and 2) the Hill equation – which we compared by τιAIC (Fig. 2C). We chose the Hill equation to model nonlinear responses due to its interpretable parameters – the empirical dose 50 (ED50) representing the SOX9 dosage at which the RE reaches half of its maximal levels, and the Hill exponent, which indicates how switch-like the RE response to SOX9 dosage change is (Fig. 2D). For all SOX9-dependent REs, the median linear – Hill τιAIC was 5.2, with a majority (73.9%) above the commonly-used cutoff of 2, indicating that most REs are better fit by the Hill equation (Fig. 2F).

In this study, we define sensitivity (or its inverse, buffering) based on the RE or gene response to decreasing SOX9 dosage from 100% to some intermediate level. Higher ED50 values (at constant Hill exponent) yield a larger response to decreases from 100% SOX9 dosage and indicate increased sensitivity, while higher Hill exponents (at constant ED50) yield a flatter response curve around 100% SOX9 dosage and decreased sensitivity. Among REs with good Hill equation fits, median values for the ED50 and Hill exponent were 19.3 and 1.99, respectively. Both values varied substantially between REs, (Fig. 2G), but the ED50 was substantially more correlated with an alternate measure of sensitivity based on the fractional change in RE accessibility when going from 100% to 50% SOX9 dosage (Extended Data Fig. 3, Spearman ρ -0.96 and -0.45 for ED50 and Hill, respectively). This indicates that the ED50, rather than the Hill exponent, is the main determinant of sensitivity/buffering in our data. Together, these results indicate a range in RE responses to SOX9 dosage changes, with most SOX9-dependent REs buffered against changes in SOX9 dosage but some showing more sensitive responses.

### Features affecting sensitivity of the RE response to SOX9 dosage changes

We next sought to identify regulatory features associated with RE sensitivity to SOX9 dosage changes. For the remainder of this paper, we use a bootstrapping approach when comparing ED50 values between groups of REs/genes to incorporate uncertainty in model fits (n=200 bootstraps, see Methods). We reasoned that the SOX9-dependent REs comprised a mix of direct effects of SOX9 binding and regulation as well as indirect, secondary effects resulting from SOX9 modulating expression of other TFs that subsequently modify chromatin accessibility. Direct SOX9 effects should arise rapidly after full SOX9 depletion in time-resolved experiments, whereas secondary effects should be delayed. We therefore performed ATAC-seq 3h after 500 nM dTAG^V^-1 treatment of *SOX9*-tagged CNCC (which we observed to result in full depletion of SOX9 within 1h, Extended Data Fig. 4A). Of the 35,713 48h SOX9-dependent REs, 9,425 showed significant (5% FDR) accessibility changes with 3h full depletion, of which almost all (96.3%) were decreases (Fig. 3A). Relative to delayed and/or upregulated REs, rapidly downregulated REs were substantially enriched for the presence of the SOX9 palindrome sequence motif (Fig. 3B), as well as SOX9 binding as assessed by V5 chromatin immunoprecipitation and sequencing (ChIP-seq) (Extended Data Fig. 4B). These results are consistent with SOX9 acting as a direct activator of REs at rapidly downregulated sites. Compared to delayed and/or upregulated REs, rapidly downregulated sites were substantially more sensitive to SOX9 dosage (Fig. 3C,D).

**Figure 3.**
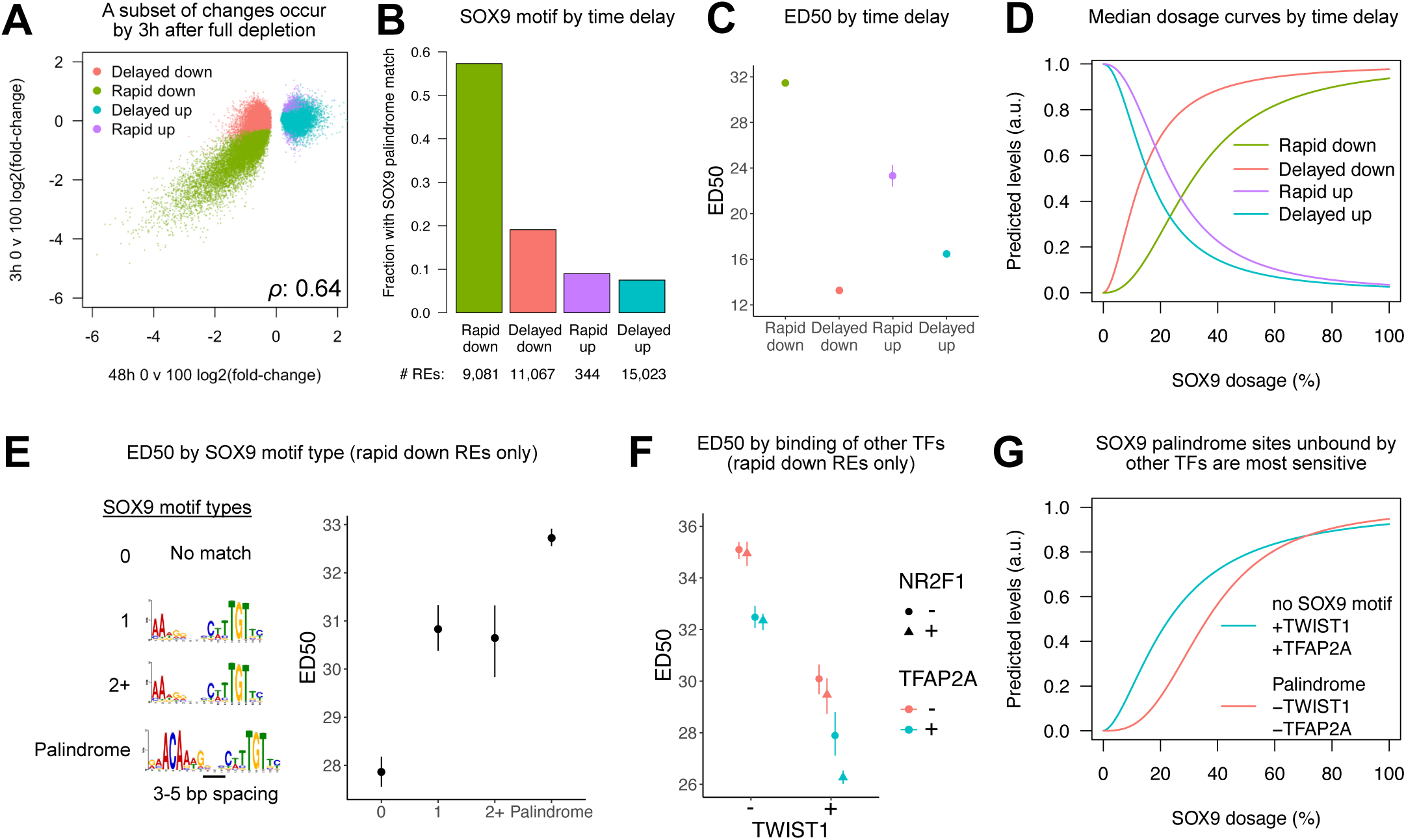
Features affecting sensitivity of the RE response to SOX9 dosage. (A) Scatterplots of full SOX9 depletion effect on chromatin accessibility at 48h (x-axis) versus 3h (y-axis) for all 48h SOX9-dependent REs. (B) Barplot indicating fraction of REs containing a SOX9 palindrome (3-5 bp spacing), stratified by time delay of response. (C) ED50 of REs stratified by time delay of response. (D) Median dosage curves based on the median fitted ED50 and Hill exponents for each group. N in (C,D) indicated in (B). (E,F) ED50 of rapid down REs stratified by either (E) SOX9 motif type, with motif position weight matrices on the left, or (F) overlap with ChIP-seq peaks for TWIST1 (x-axis) TFAP2A (color), and NR2F1 (shape). N of groups from left to right in (E): 2,263, 1,315, 360, 5,221. N of groups from left to right in (F): 1,874, 736, 985, 1,805, 774, 602, 296, 2,087. (G) Median dosage curves as in (D) for the indicated combinations of SOX9 motif and binding of other TFs. Points and error bars in (C,E,F) represent median and 95% confidence intervals as computed by bootstrap (see Methods).

We focused on direct SOX9 target sites, seeking to identify additional features associated with sensitivity among these REs. SOX9 has been reported to bind different types of motifs, including single SOX9 elements, partial palindromes, and full palindromes with a 3-5 bp spacing. We found that REs containing the full SOX9 palindrome with 3-5 bp spacing were more sensitive than sites containing either one or multiple partial palindromes, with REs containing no detected motif being the least sensitive (Fig. 3E). The SOX9 palindrome with 3-5 bp spacing was also specifically associated with a modest increase in the Hill exponent, consistent with previous reports showing its requirement for cooperative SOX9 binding (Extended Data Fig. 4C,D)^41^. Thus, SOX9-intrinsic features (the type of motif and resultant mode of SOX9 binding) further modulate RE sensitivity to SOX9 dosage changes among its direct targets. Indeed, direct targets with larger effects of SOX9 depletion were most sensitive to SOX9 dosage changes (Extended Data Fig. 5A).

Finally, we assessed whether SOX9-extrinsic factors could also modulate RE sensitivity to SOX9 dosage. We focused on binding by other TFs, specifically TWIST1, NR2F1, and TFAP2A, as they have well-known roles in CNCCs and their binding in hESC-derived CNCCs has previously been characterized by ChIP-seq^32, 38^. We found that increased binding of other TFs substantially decreased RE sensitivity to SOX9 dosage; the strongest, independent effects were seen for TWIST1 and TFAP2A binding, with minor effects of NR2F1 binding at REs bound by both TWIST1 and TFAP2A (Fig. 3F). We replicated this result when stratifying REs only based on corresponding TF sequence motifs (Extended Data Fig. 5B). Baseline levels of both the active histone mark H3K27ac and chromatin accessibility were negatively correlated with sensitivity, with a stronger effect of chromatin accessibility (Extended Data Fig. 5C,D). Combining SOX9 motif type and binding by other TFs revealed that REs containing the SOX9 palindrome motif that were also unbound by other TFs showed the most sensitivity to SOX9 dosage (Fig. 3G, Extended Data Fig. 5E). Together, these results indicate that a combination of SOX9-intrinsic and -extrinsic factors dictate the relative sensitivity or buffering of SOX9-dependent REs to SOX9 dosage. REs directly bound and strongly regulated by SOX9 where SOX9 is likely the primary TF driving accessibility are most sensitive, whereas REs indirectly regulated by SOX9 through secondary effects and/or to which other TFs bind to and promote accessibility are most buffered.

### The chromatin accessibility response at highly contributing REs predicts cognate gene responses to SOX9 dosage changes

We next assessed the gene expression response to SOX9 dosage by analzying RNA-seq on the same SOX9 dosage series. Individual genes showed diverse responses (Fig. 4A, Extended Data Fig. 6A). Changes in SOX9 dosage had overall nonlinear effects on gene expression in principal component space (Extended Data Fig. 6B), and most (70.3%) of the 1,232 SOX9-dependent genes (of which 688 decreased and 544 increased upon full depletion, 5% FDR) were better fit by a Hill than a linear equation (Fig. 4B) (Supplementary Table 2), with variability in the ED50 and Hill exponents (Fig. 4C). Thus, like REs, most genes are buffered against SOX9 dosage, with a subset more sensitive.

**Figure 4.**
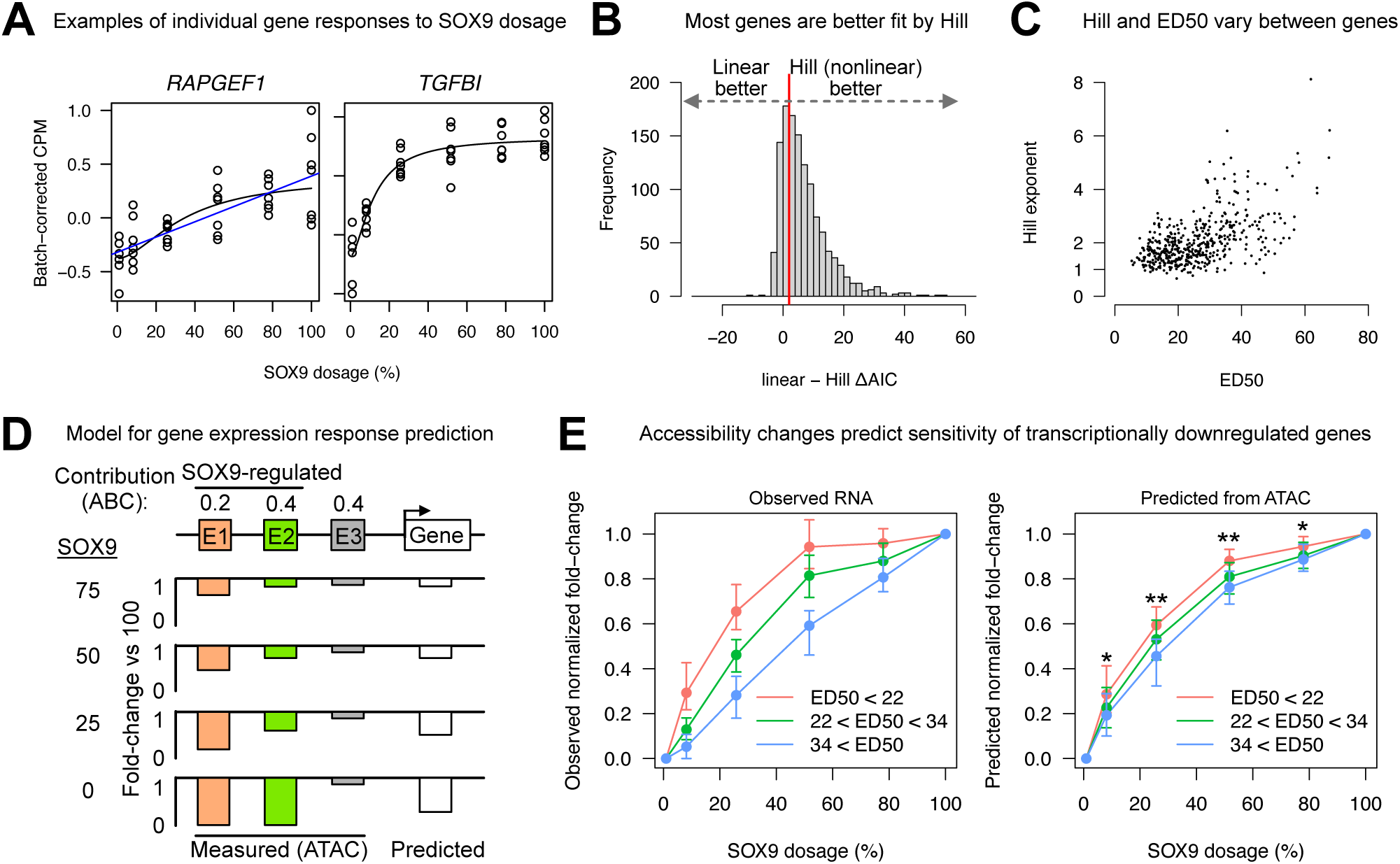
RE dose-response curves predict the shape of gene dose-response curves. (A) Examples of genes with sensitive (left) or buffered (right) responses to SOX9 dosage changes, as assessed by RNA-seq. Black and blue lines represent Hill and linear fits, respectively. (B) Histogram of ι1AIC of all SOX9-dependent genes, calculated as in Fig. 2C. Red line indicates ι1AIC = 2. (C) ED50 and Hill exponent across individual SOX9-dependent genes with good fit (p < 0.05 for both parameters). (D) Schematic of approach to predict RNA level changes based on Activity-by-Contact (ABC) contribution scores and fold-changes of REs at each SOX9 dosage. (F) Distributions of observed (left) or predicted (right) fold-changes vs full SOX9 dosage at each concentration, normalized to full depletion fold-change and stratified by ED50 (colors). Only genes transcriptionally downregulated by 24h full SOX9 depletion (assessed by metabolic labeling, SLAM-seq) are analyzed. N of groups by color: red, 66; grey, 77; blue, 57. Points and error bars represent median and 25^th^ and 75^th^ percentiles of distribution. * p < 0.01, ** p < 1e-6.

Given the overall similar responses of chromatin accessibility and gene expression to SOX9 dosage, we investigated whether RE responses predict the responses of their cognate target genes. We focused our predictions on the subset of SOX9-dependent genes that also show changes in nascent transcription in response to SOX9 depletion. We performed Thiol (SH)-linked alkylation for the metabolic sequencing of RNA (SLAM-seq)^42^ to quantify changes in newly transcribed mRNA levels after either 3h or 24h of full SOX9 depletion, finding that of the 1,232 48h SOX9-dependent genes, 122 (62 down, 60 up) responded significantly (10% FDR) at 3h, and 395 (206 down, 189 up) responded at 24h. Effect sizes at 24h were correlated with and generally larger in magnitude than those at 3h (Extended Data Fig. 6C), and known direct SOX9 targets such as *COL2A1*^43^ responded at 24h but not 3h (Extended Data Fig. 6D), suggesting a time lag between the effects of SOX9 depletion on chromatin and transcription. Accordingly, we sought to predict the gene expression responses to SOX9 dosage for the larger set of SOX9-dependent genes that responded transcriptionally to 24h of SOX9 depletion.

The Activity-by-Contact (ABC) model has been recently shown to effectively predict RE-gene connections by computing each RE’s contribution to gene transcription (ABC score) as its ‘Activity’ (a combination of accessibility and the active histone mark H3K27ac) divided by its contact (estimated by chromatin conformation capture or a genomic distance-power law function), normalized to the contributions of all other REs within 5 Mb of the TSS^44^. We reasoned that the ABC model could be extended to predict the effect of multiple REs changing in ‘Activity’ at each SOX9 concentration relative to full SOX9 dosage. Although the ‘Activity’ portion of ABC includes H3K27ac levels, we observed that the effect of full SOX9 depletion on chromatin accessibility and H3K27ac was highly correlated (Extended Data Fig. 6E,F). In effect, the fold-change in a gene’s expression at a reduced SOX9 dosage relative to full dosage is predicted as the average fold-change in accessibility at all REs within 5Mb of the TSS, weighted by each RE’s contribution, or ABC score (Fig. 4D).

We first assessed whether we could predict the direction of gene expression change in response to SOX9 dosage. We stratified transcriptionally regulated genes (responding by SLAM-seq after 24h SOX9 full depletion) into those increasing or decreasing in response to SOX9 dosage and plotted their fold-changes at each SOX9 dosage level vs 100 along with genes not responding at all to SOX9 depletion (Extended Data Fig. 6G, left). The predicted fold-changes for these genes clearly and significantly stratified them in the same manner, with predictions for downregulated genes performing substantially better than predictions for upregulated genes (Extended Data Fig. 6G, right).

Given that predictions for genes transcriptionally downregulated in response to SOX9 depletion were the most accurate, we next focused on predicting differences in SOX9 dosage sensitivity among this set of genes. We grouped these genes into three bins based on their observed sensitivity to SOX9 (defined as specific ranges of ED50 values), normalizing the fold-change at each SOX9 dosage to the fold-change at full SOX9 depletion (Fig. 4E, left). Normalized fold-changes based on RE predictions clearly separated the three groups (Fig. 4E, right), although to a lesser extent than the directionality predictions. Inspection of top predictions indicated that our simple model could predict either buffered (*TENT5B*) or sensitive (*SOX5*) responses (Extended Data Fig. 6H). Together, these results indicate that broadly, the nature of the RE response to SOX9 dosage changes (sensitive or buffered) translates into the response of cognate genes based on the RE’s overall contribution to that gene’s transcription.

### The pro-chondrogenic program is sensitized to SOX9 dosage changes

Our results thus far indicate that while most SOX9-dependent REs are buffered against small to moderate changes in SOX9 dosage, a subset is more sensitive due to certain regulatory features, and that SOX9- sensitive RE responses often translate into gene expression responses. We next sought to assess the impact of such dosage-sensitive genes on cellular phenotypes, focusing on chondrogenic differentiation potential, as SOX9 has well-known roles in entry into the chondrogenic program and chondrogenesis itself (Lefebvre and Dvir-Ginzberg, 2017). We first tested whether pro-chondrogenic genes showed heightened sensitivity to SOX9 dosage changes in CNCCs. Genes both annotated to play a role in cartilage development and upregulated during *in vitro* chondrogenesis (“pro-chondrogenic genes”) showed substantially higher ED50 values than other groups of genes (Fig. 5A). Examples of sensitive pro-chondrogenic genes with well-known roles include the collagen-encoding genes *COL11A1* and *COL2A1*, as well as genes encoding other transcriptional regulators such as *SOX5* (Fig. 5B). We did not observe a similar pattern of sensitivity when stratifying genes upregulated in response to SOX9 depletion in the same way (Extended Data Fig. 7A), suggesting that the pro-chondrogenic functions of SOX9 may be especially dosage sensitive.

**Figure 5.**
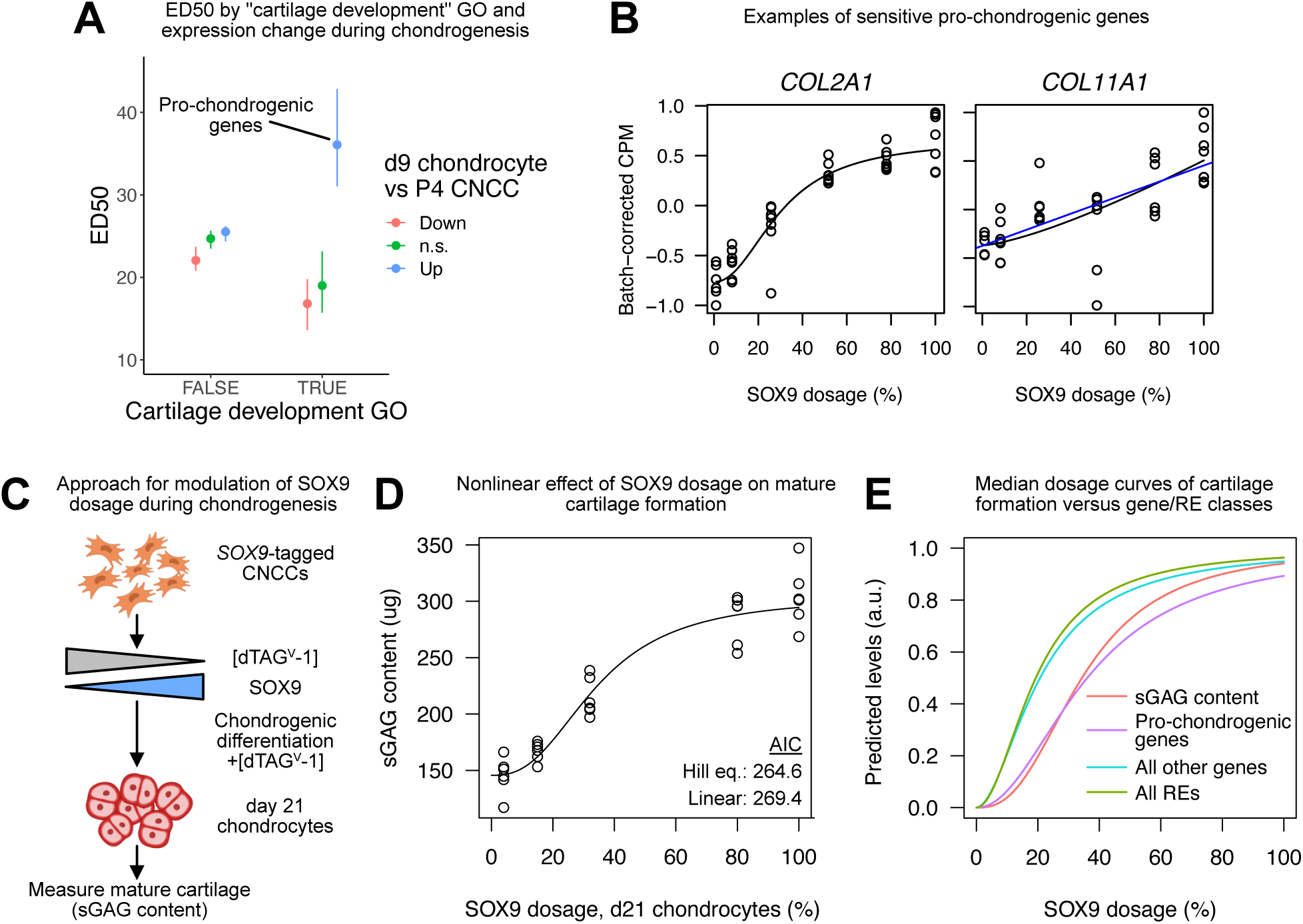
The pro-chondrogenic program is sensitive to SOX9 dosage. (A) ED50 of SOX9-downregulated genes stratified by presence in the “Cartilage development” Gene Ontology (GO) category (x-axis), and expression change in chondrocytes compared to CNCCs (color, data from Long et al 2020). N of groups from left to right: 251, 486, 445, 9, 13, 28. Points and error bars in represent median and 95% confidence intervals as computed by bootstrap (see Methods). (B) Examples of known pro-chondrogenic genes that are sensitive to SOX9 dosage. (C) Schematic of approach to titrate SOX9 dosage during 21-day chondrogenic differentiation. (D) Sulfated glycosaminoglycan (sGAG, representative of mature cartilage) at day 21 of chondrogenesis as a function SOX9 dosage as estimated in Extended Data Fig. 7B. Black curve represents Hill equation fit. (E) Median dosage curves based on fitted ED50 and Hill exponents for all REs and genes, pro-chondrogenic genes (purple, labeled group in (A), and sGAG content (from (D)).

To test whether increased sensitivity of pro-chondrogenic genes results in increased sensitivity of chondrogenesis, we modulated SOX9 dosage during differentiation of CNCCs to chondrocytes *in vitro*. We treated *SOX9*-tagged CNCCs with differing dTAG^V^-1 concentrations and then performed 21-day chondrogenic differentiations using established protocols^32^, maintaining the same dTAG^V^-1 concentrations throughout (Fig. 5C). Quantification of SOX9 levels by flow cytometry indicated the ability to precisely modulate SOX9 dosage to five distinct levels throughout chondrogenesis (Extended Data Fig. 7B). To quantify the degree of resulting chondrogenesis, we measured total levels of sulfated glycosaminoglycans (sGAGs), linear polysaccharides that mark the extracellular matrix of mature cartilage, using a colorimetric assay based on the binding of sGAGs to 1,9-dimethylmethylene blue. This revealed a nonlinear relationship between SOX9 dosage and functional chondrogenesis as measured by sGAG content (Fig. 5D). There was no effect of the highest dTAG^V^- 1 concentration (500 nM) on WT CNCC chondrogenesis, indicating minimal off-target effects of dTAG^V^-1 on chondrogenesis (Extended Data Fig. 7C). Importantly, the fitted SOX9 dosage-sGAG curve more closely matched the curve for pro-chondrogenic genes than other genes or REs (Fig. 5E). Thus, *in vitro* chondrogenesis is sensitized to SOX9 dosage, more so than most genes or REs, likely due to the heightened sensitivity of important pro-chondrogenic genes.

### Genes associated with PRS-like phenotypes are sensitized to SOX9 dosage changes

We next assessed the impact of SOX9-sensitive genes and REs on human morphological and craniofacial disease phenotypes. We first examined known craniofacial disorder associations of genes that are downregulated upon SOX9 depletion, also considering the relation to PRS, which is caused by heterozygous LoF coding or noncoding mutations at the *SOX9* locus. However, mutations in genes other than *SOX9* have also been associated with PRS-like micrognathia phenotypes in either humans or mice (reviewed in ^46^).

We observed that SOX9-dependent genes associated with dominant (likely more dosage-sensitive) craniofacial disorders unrelated to PRS had lower ED50 values than genes not associated with craniofacial disorders, while genes associated with recessive (likely less dosage-sensitive) disorders had higher ED50 values (Fig. 6A,B); this suggests buffering of important, dosage-sensitive genes that strongly impact craniofacial development but are nevertheless ultimately SOX9-dependent. However, genes associated with PRS-like craniofacial defects (Motch Perrine et al., 2020) showed higher ED50 values than either group, and were thus most sensitive to SOX9 dosage changes (Fig. 6A,B). Genes in this group include *COL2A1* and *COL11A1*, of which haploinsufficiency is associated with subtypes of Stickler syndrome^47, 48^, which like PRS, includes underdevelopment of the lower jaw, and which are also associated with pro-chondrogenic functions as described above. Similar results were not observed when only considering genes that are upregulated upon SOX9 depletion (Extended Data Fig. 8). Thus, while dosage-sensitive, SOX9-dependent craniofacial genes are generally buffered against changes in SOX9 dosage, those with PRS-like phenotypes are highly sensitive.

**Figure 6.**
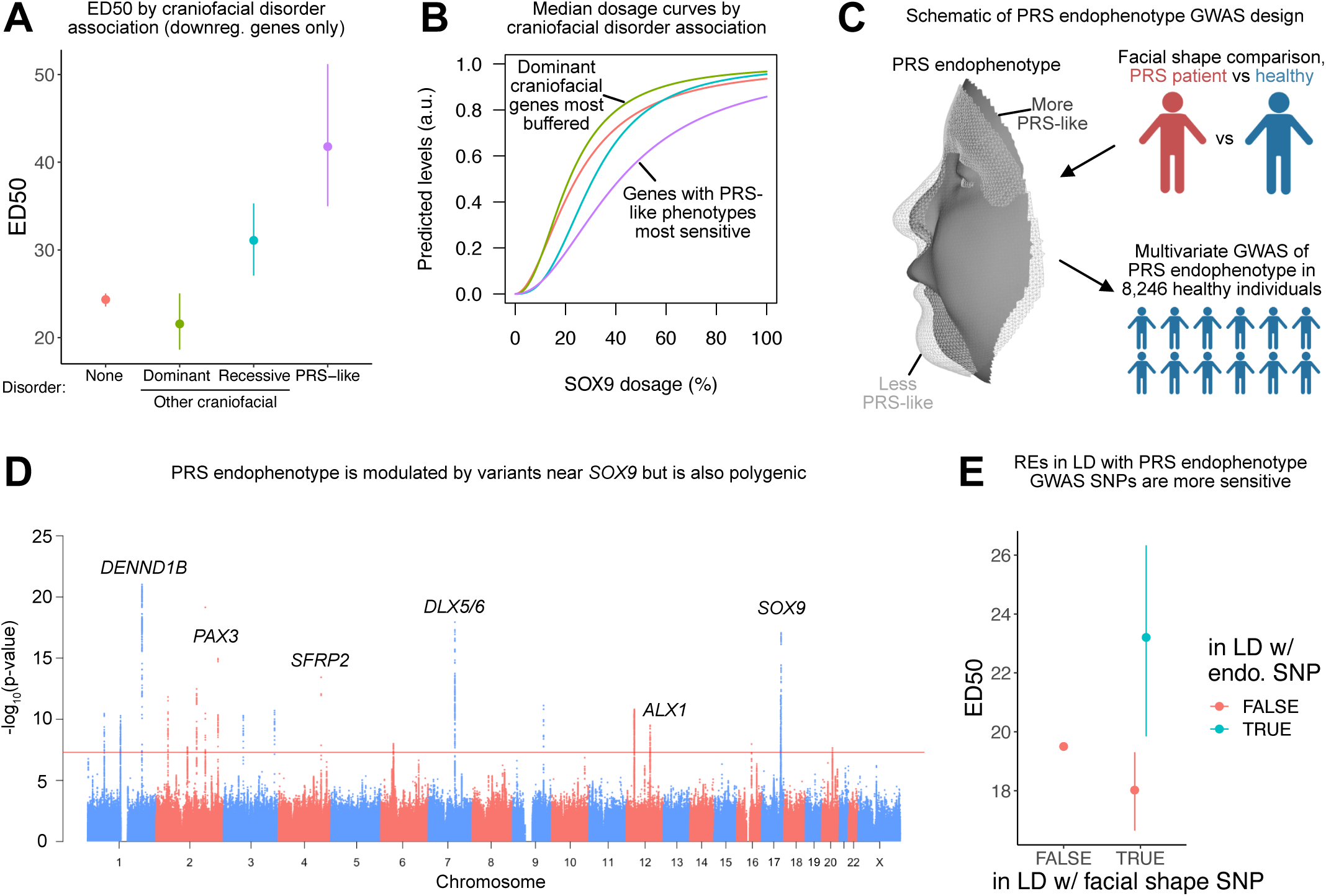
Genes and REs associated with Pierre Robin Sequence (PRS)-like phenotypes are uniquely sensitized to SOX9 dosage. ED50 (A) of SOX9-downregulated genes stratified by craniofacial disorder association. PRS-like associations as defined by Perrine et al 2020. N of groups from left to right: 665, 13, 6, 8. (B) Median dosage curves (based on both ED50 and Hill exponent) of the same groups from (A). (C) Schematic of approach to conduct a multivariate GWAS on a PRS-defined endophenotype in healthy human individuals. (D) Manhattan plot of PRS endophenotype GWAS. Black line indicates genome-wide significance. Candidate genes near top GWAS signals are labeled. (E) ED50 of REs stratified by linkage disequlibrium (LD, r^2^ > 0.5) with any facial shape GWAS lead SNP (x-axis, as defined in Naqvi, Hoskens et al. 2022) and further with any lead SNP associated with the PRS endophenotype GWAS from (D) (color). N of groups from left to right: 35,450, 209, 54. Points and error bars in (A,E) represent median and 95% confidence intervals as computed by bootstrap (see Methods).

These observations suggest that phenotypic specificity associated with SOX9 dosage perturbation during craniofacial development is mediated by a subset of genes sensitized to changes in SOX9 dosage that includes key pro-chondrogenic effector genes.

### GWAS of PRS endophenotype in healthy individuals uncovers selective sensitivity of linked REs to SOX9 dosage changes

We next assessed whether similar principles, as described above for gene-disorder associations, apply to common variation in facial shape in healthy individuals. We sought to first identify SNPs associated with normal-range variation in healthy individuals along the facial shape axis from typical to PRS (PRS endophenotype). We compared three-dimensional facial scans from 8,246 healthy individuals and 13 PRS patients to define a univariate PRS endophenotype (Extended Data Fig. 9A) for which each individual could be scored in each of 63 facial segments obtained by hierarchical spectral clustering (Extended Data Fig. 9B)^29^. We then tested which facial segments showed significant (nominal *p* < 0.05) differences in the endophenotype score between PRS patients and the 8,246 healthy European-ancestry individuals (Extended Data Fig. 9C,D). From these 30 facial segments, we conducted a multivariate GWAS for the PRS endophenotype in the healthy, European-ancestry individuals, combining the endophenotype scores from each segment using canonical correlation analysis^29^ (Fig. 6C, Extended Data Fig. 9E,F).

We observed two independent GWAS signals at the *SOX9* locus (Extended Data Fig. 10), but 20 additional loci across the genome reached genome-wide significance (*p* < 5 x 10^-8^) (Supplementary Table 3). This indicates that phenotypic variation along the healthy-to-PRS axis is modulated by variants near *SOX9*, as would be expected given associations between *SOX9* mutations and PRS itself, but is also polygenic (Fig. 6D). The 20 genome-wide significant loci were a subset of loci identified by previous studies; we thus segregated previously reported facial GWAS lead SNPs based on significant association with the PRS endophenotype (Bonferroni-corrected *p* < 0.05). SOX9-dependent REs in linkage disequilibrium (LD, r^2^ > 0.5) with signals for facial shape phenotypes not associated with the PRS endophenotype had slightly lower ED50 values than other SOX9-dependent REs. In contrast, facial shape-linked REs in LD with PRS endophenotype signals had higher ED50 values (Fig. 6E). Combined with the above analyses of craniofacial disorder-associated genes, these results indicate that REs and genes with phenotypic impact on normal-range and disease-associated variation unrelated to that of SOX9 are generally buffered against changes in TF dosage, even if they ultimately are SOX9-dependent, while REs and genes with similar (PRS-like) phenotypes are most sensitive.

## DISCUSSION

Here, we have quantified the relationship between TF dosage and facial phenotype at multiple biological levels – molecular, cellular, and morphological – using SOX9 as a model. Starting with effects on chromatin, we observed that most SOX9-dependent REs are buffered in their chromatin accessibility response, with limited changes observed until low (∼20% or less) SOX9 dosage. However, a subset is more sensitive to SOX9 dosage and is enriched for specific regulatory features, including direct binding and regulation by SOX9 but lack of binding by other important TFs. Such sensitive REs lead to gene expression responses to SOX9 that are also more sensitive than most. Among SOX9-dependent genes, a set of pro-chondrogenic genes explains the heightened sensitivity of *in vitro* chondrogenesis to SOX9 dosage. Finally, genes and REs with similar phenotypic associations as SOX9 are sensitive to SOX9 dosage, in contrast to other key craniofacial genes and REs with distinct phenotypic associations from SOX9 that are generally buffered.

We propose a model of the regulatory network downstream of SOX9 that synthesizes these observations (Fig. 7A). In this model, REs regulated by SOX9 exist on a scale from sensitive to buffered as a result of the combination of regulatory features discussed above. Genes with nearby sensitive REs will themselves show more sensitive responses to SOX9 dosage, while those with highly contributing and buffered REs are more robust to perturbation yet are still ultimately SOX9-dependent. This range of gene and RE responses (from sensitive to buffered) intersects nonrandomly with phenotypic impact - those with generally important roles in CNCC biology but causing phenotypes distinct from SOX9 are buffered against SOX9 dosage change, but a subset of sensitive genes corresponds to known, dosage-sensitive effectors of specific cellular processes, which in turn are important for aspects of morphology most similar to those of SOX9. Specifically, our results demonstrate that several key pro-chondrogenic genes and *in vitro* chondrogenesis are sensitized to SOX9 dosage changes. Thus, the observed sensitivity of key chondrogenesis effector genes and of chondrogenesis itself to SOX9 dosage changes could account for the specificity of SOX9-associated PRS-like mandibular phenotypes, perhaps involving Meckel’s cartilage, a cartilage ’template’ that is specifically involved in formation of the mandible^49^.

**Figure 7.**
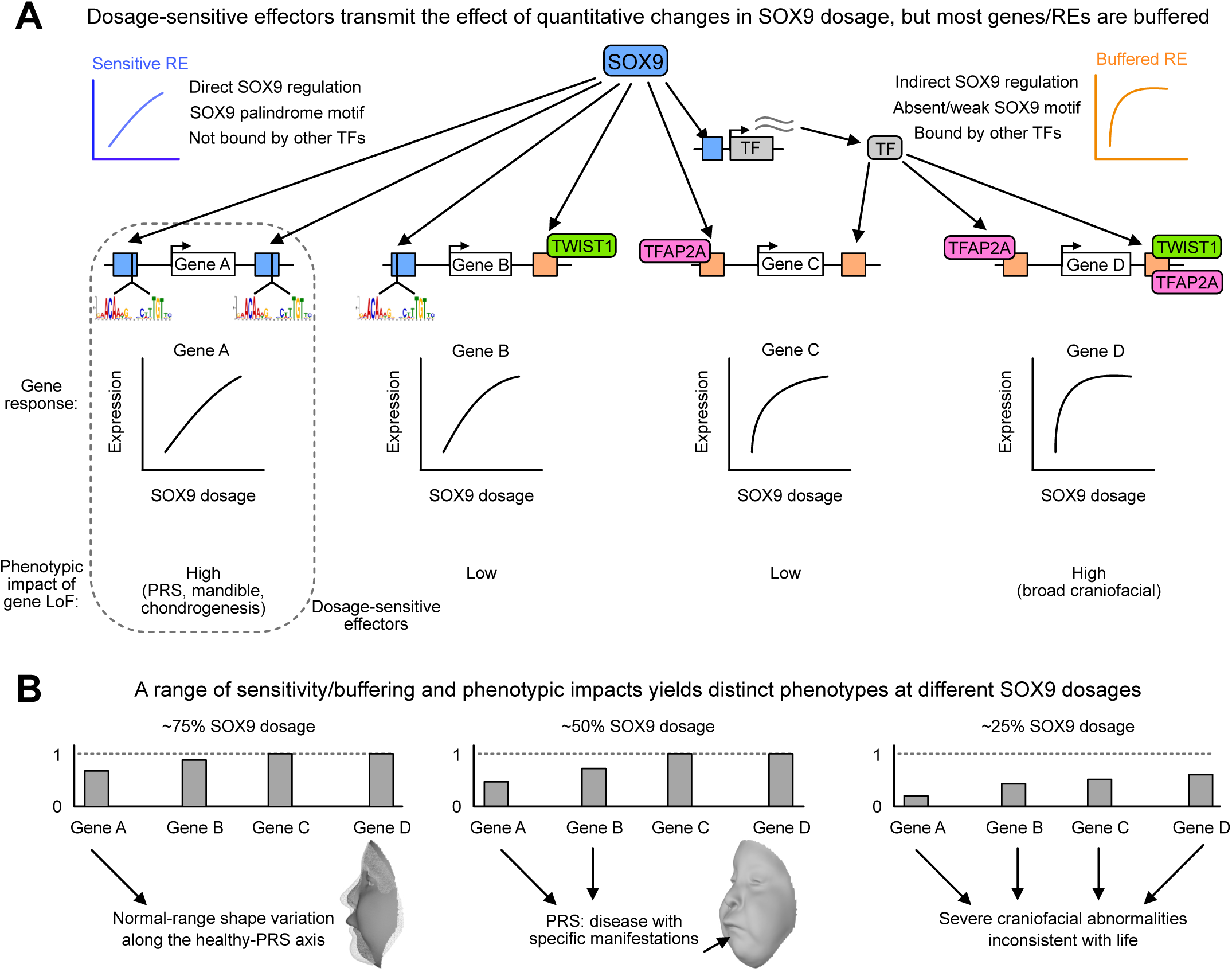
Dosage-sensitive effectors transmit the effect of quantitative changes in SOX9 dosage to provide phenotypic specificity. (A) Schematic indicating which features make REs sensitive (blue) or buffered (orange) to SOX9 dosage. Dosage-sensitive effectors are REs and genes mediating changes in cellular behaviors as a result of quantitative changes to their activity or expression, and thus with high phenotypic impact (dotted box), as compared to sensitive genes with low phenotypic impact (Gene B) or phenotypically impactful genes that are buffered (Gene D). (B) Schematic of gene expression changes in response to the indicated SOX9 dosages based on sensitivities from (A). Arrows indicate gene’s contribution to phenotype. PRS, Pierre Robin Sequence; LoF, loss of function.

Our model can explain distinct phenotypes observed across the full range of SOX9 dosage (Fig. 7B). In the ∼80-100% dosage regime associated with common genetic variation at the *SOX9* locus, most downstream effects at dosage-sensitive genes affecting chondrogenesis and mandibular development are subtle, resulting in common phenotypic variation along the healthy-PRS axis. At dosages closer to ∼50%, the further decreased activity of dosage-sensitive effector genes (and potentially, other effects contributed by additional genes affected at this dosage) exacerbates the phenotypic effects, resulting in the disease phenotype of PRS itself. Finally, even lower SOX9 dosages (∼25% or less) lead to a broader dysregulation of other genes and cellular processes important for craniofacial development; these changes, combined with further increased perturbations to the dosage-sensitive effectors, result in wide phenotypic impacts and embryonic lethality^50^.

Several lines of evidence suggest that the core concepts of our model – a range of TF-dependent REs and genes from sensitive to buffered combined nonrandomly with a range of phenotypic importance – also apply to other craniofacial TFs, and more broadly to other traits. Haploinsufficiency of many other craniofacial TFs is often associated with syndromes comprising characteristic facial features (*e.g*. PAX3 in Waardenburg, TWIST1 in Saethre-Chotzen, TFAP2A in Branchiooculofacial syndromes, respectively), but similar to SOX9, these TFs bind to and presumably regulate thousands of REs (and perhaps hundreds of genes) across the genome. Groups of effector genes uniquely sensitive to the dosage of each TF would result in phenotypic specificity at ∼50% TF dosage found in haploinsufficient syndromes, while most TF-regulated genes having buffered responses would still allow for widespread binding and regulation. A recent study assessed dose-dependent effects of the cardiac TF TBX5 on gene expression in iPSC-derived cardiomyocytes, finding that a subset of genes dysregulated by homozygous *TBX5* deletion are also dysregulated, but to a lesser extent, by heterozygous deletion; some of these genes may represent dosage-sensitive effectors for *TBX5*^51^.

Our model simultaneously allows for both robustness and phenotypic sensitivity to TF dosage. Robustness can be explained by nonlinear relationships between gene dosage and phenotype, which are suggested by human and mouse genetics. The vast majority of GWAS variants, which typically have small effects on expression and thus sample the dosage curve close to 100% dosage, have additive, linear effects on phenotype^52, 53^, whereas large-effect rare variants associated with Mendelian disease show nonlinearity in phenotypic effects as indicated by recessivity (most diseases with dominant inheritance can also be thought as recessive with respect to lethality). Experimental manipulation of *Fgf8* dosage in mice followed by quantification of craniofacial morphology^54^ indicates a similarly nonlinear dosage-to-phenotype relationship. Our study, finding overall nonlinear effects of SOX9 dosage, is consistent with these findings. However, our model suggests that these overall nonlinear effects are a composite of distinct molecular responses: most SOX9-regulated REs and genes are buffered against moderate changes in SOX9 dosage, while trait variation and disease is caused by the SOX9-sensitive effectors that show substantial effects at ∼50% SOX9 dosage. Buffered REs and targets can explain robustness to TF dosage perturbation, while the latter class of sensitive effectors mediate phenotypic specificity associated with TF dosage changes.

How do TFs act to modulate variation in complex trait and common disease risk, most of which is highly polygenic^55^? One possibility is by regulating broad transcriptional programs, in which the downstream effects of a trait-associated TF are distributed widely among many targets genome-wide. While SOX9, like many TFs, regulates thousands of RE/gene targets, our study indicates most of these targets are buffered and have individually tiny effects at the < 50% variation in TF dosage observed in GWAS. These minimal effects can contribute substantially to variation when summed up genome-wide, but most effects with individually appreciable contributions to variation result from a subset of REs and genes sensitive to SOX9 dosage changes. Among the sensitive genes, even a smaller subgroup represents phenotypic effector genes of chondrogenesis and PRS-like phenotypes. Such effector genes are conceptually similar to core genes in the omnigenic model^56, 57^. Core genes act directly on a trait, but their expression can be affected by perturbations to many peripheral genes, but especially master regulators (such as TFs) that can modulate expression of multiple core genes simultaneously. The range of gene responses from sensitive to buffered that we have found may represent a way of limiting the effects of master regulators to core genes at the subtle variation in TF dosage most relevant in GWAS.

## ACKNOWLEDGEMENTS

We thank members of the Wysocka and Pritchard labs, as well as James Ferrell, for helpful discussions and comments. This research was supported by NIH grant R01HG008140 and funding from the Howard Hughes Medical Institute. S.N. was supported by a Helen Hay Whitney Fellowship. S.K. was supported by a Damon Runyon Fellowship. JW was supported by a Lorry Lokey endowed professorship and a Stinehart Reed award. The KU Leuven research team and analyses were supported by the National Institutes of Health (1-R01-DE027023, 2-R01-DE027023), The Research Fund KU Leuven (BOF-C1, C14/15/081 & C14/20/081) and The Research Program of the Research Foundation - Flanders (Belgium) (FWO, G078518N). RS, BH and OK were supported by NIH U01DE024440. The computational resources and services used in this work were provided by the VSC (Flemish Supercomputer Center), funded by the Research Foundation - Flanders (FWO) and the Flemish Government – department EWI. Some of the computing for this project was performed on the Sherlock cluster. We thank Stanford University and the Stanford Research Computing Center for providing computational resources and support that contributed to these research results.

## AUTHOR CONTRIBUTIONS

Conceptualization, S.N., T.S., J.K.P, and J.W.;

Formal Analysis, S.N., S.K., H.H., and H.S.M.;

Investigation, S.N. and S.K.;

Resources, R.A.S, O.D.K, P.C., J.K.P, and J.W.;

Writing - Original Draft, S.N.;

Writing - Review & Editing, S.N., J.K.P, and J.W.;

Visualization, S.N., H.H., and H.S.M.;

Supervision, J.K.P and J.W.;

Funding Acquisition, J.K.P and J.W.

## RESOURCE AVAILABILITY

### Materials availability

Plasmids generated in this study will be deposited in Addgene by the date of publication.

### Data availability

The raw sequencing files generated during this study will be available on GEO (accession number GSE205904); corresponding processed data is available on Zenodo (10.5281/zenodo.6596465). Data availability for the 8,246 healthy individuals used for PRS endophenotype GWAS are described in ^30^. Facial scans from PRS patients (used to define the PRS endophenotype) are available through the FaceBase Consortium (https://www.facebase.org FB00000861) under controlled access.

### Code availability

Custom code used for analysis of processed sequencing data is available on Zenodo (10.5281/zenodo.6596465). KU Leuven provides the MeshMonk (v.0.0.6) spatially dense facial-mapping software, free to use for academic purposes (https://github.com/TheWebMonks/meshmonk). Matlab 2017b implementations of the hierarchical spectral clustering to obtain facial segmentations are available from a previous publication^58^.

## ONLINE METHODS

### Cell culture

Female H9 (WA09; RRID: CVCL_9773) human embryonic stem cells (hESCs) were obtained from ATCC and cultured in either mTeSR1 (Stem Cell Technologies) for at least one passage prior to differentiation into cranial neural crest cells (CNCCs) or mTeSR Plus (Stem Cell Technologies) for gene editing, single-cell cloning, expansion, and maintenance. hESCs were grown on Matrigel Growth Factor Reduced (GFR) Basement Membrane Matrix (Corning) at 37°C. hESCs were fed every day for mTeSR1 or every two days for mTeSR Plus and passaged every 5-6 days using ReLeSR (Stem Cell Technologies).

HEK293FT cells were obtained from ATCC and cultured in complete media. Complete media: DMEM-HG, 10% FBS, 1x Non-essential amino acids, 1x Glutamax, 1x antibiotic/antimycotic. Cells were fed every other day and passaged every 2-3 days using trypsin.

### Adeno-associated virus (AAV) production and titration for CRISPR/Cas9 genome editing

Production of AAV for genome-editing was produced as described previously ^36^, with the following modifications. Left and right homology arms (∼1kb) surrounding the *SOX9* stop codon, flanking the linker-FKBP12FV36-linker-mNeonGreen-linker-V5-stop tag, were cloned into an AAV backbone. This vector plasmid, along with the AAV6 packaging plasmid pDGM6 (Addgene), were transfected into 70-80% confluent, early-passage HEK293FT cells seeded 24 hours prior to transfection at ∼8-9 million cells per 15cm plate, and changed with fresh media 2-6 hours before transfection. For each 15 cm plate (two per individual AAV6 prep), the transfection mix was: 22 ug pDGM6, 6 ug vector plasmid, 120g Polyethyenimine, and Opti-MEM to 1mL. Cells were changed into slow-growth media 24 hours post-transfection. Slow-growth media: Same as complete media but with 1% FBS. Cells were harvested 48 hours after changing to slow-growth media with the AAVpro Purification Kit Midi (Takara, 6675) as per manufacturer instructions.

Titration of purified AAV6 was done by qPCR. Briefly, a previously flash-frozen and thawed 10uL aliquot of virus was treated with TURBO DNase (Invitrogen, AM2238) as per manufacturer instructions to digest unpackaged DNA. DNAse was inactivated by 0.001M EDTA (final concentration) and incubation at 75°C for 10 min. Virus DNA was released by proteinase K treatment at 50°C for two hours to overnight. Proteinase solution: 1M NaCl, 1% w/v N-lauroylsarcosine, 100 ug/mL Proteinase K (Invitrogen, 25530049). Samples were then boiled for 10 min, and diluted twice in H20 such that the final dilution was 1:200,000. DNA standards comprising 10^10^-10^3^ molecules were prepared using inverted terminal repeat (ITR)-containing AAV6 backbone plasmids. qPCR was performed on standards and test samples using the Lightcycler 480 Probes Master kit (Roche, 04707494001) with the following ITR probe and primer sequences: probe, 5’-FAM-CACTCCCTCTCTGCGCGCTCG-BBQ; ITR-F, GGAACCCCTAGTGATGGAGTT, ITR-R, CGGCCTCAGTGAGCGA.

### Generation of CRISPR/Cas9 and AAV genome-edited cell lines

Genome editing of hESCs was performed as previously described ^36^, with the following modifications. hESCs were pre-treated with 10uM RHO/ROCK pathway inhibitor Y-27632 (Stem Cell Technologies, 72304) for 2-24 hours, harvested and brought to single cells with accutase and vigorous pipetting, and ∼800,000 nucleofected with a Cas9-sgRNA ribonucleoprotein (RNP) complex using the Lonza 4D Nucleofection system. RNP consisted of 17 ug Sp-Cas9 HiFi (IDT) and 300 pmol sgRNA duplex. Cells were plated on Matrigel-coated plates with mTeSR Plus with 10 uM Y-27632 and AAV at desired MOI (typically ∼25,000). Cells were changed into mTeSR Plus with 10 uM Y-27632 but no AAV 4-24 hours after initial plating, and an additional equal volume of mTeSR Plus with no Y-27632 was added 2 days later. Subsequent feedings were done with no Y-27632 until cells approached confluency, at which point cells were again harvested and brought to single cells with accutase (after 10 uM Y-27632 pre-treatment) and 500-1000 cells were plated per well of a 6-well plate. Cells were then expanded until colonies were of sufficient size to pick, before which cells were again pre-treated with 10 uM Y-27632 for 2-24 hours. Colonies were picked into 24- or 48-well plates without Y-27632 and allowed to expand for ∼5 days and passaged 1:2 using ReLeSR, with one half plated on another 24- or 48- well plate and the other half used for lysis with QuickExtract (Lucigen, QE9050). Genotyping PCR was performed with one primer outside the homology arms and one primer inside the opposite homology arm. Clones containing the desired knock-in were expanded and used for genomic DNA extraction with the Quick-DNA miniprep kit (Zymo D3024), followed by the same genotyping PCR and Sanger sequencing to confirm knock-in.

### Differentiation of hESCs to CNCCs and chondrocytes

hESCs were differentiated to cranial neural crest cells (CNCCs) using a protocol described previously ^37, 38^. Briefly, hESCs were grown for 5-6 days until large colonies formed, then were disaggregated using collagenase IV and gentle pipetting. Clumps of ∼200 hESCs were washed in PBS and transferred to a 10cm Petri dish in neural crest differentiation media (NDM). NDM: 1:1 ratio of DMEM-F12 and Neurobasal, 0.5x Gem21 NeuroPlex Supplement With Vitamin A (Gemini, 400-160), 0.5x N2 NeuroPlex Supplement (Gemini, 400-163), 1x antibiotic/antimycotic, 0.5x Glutamax, 20ng/ml bFGF (PeproTech, 100-18B), 20ng/ml EGF (PeproTech, AF-100-15) and 5ug/ml bovine insulin (Gemini Bio-Products, 700-112P). After 7-8 days, neural crest emerged from neural spheres attached to the Petri dish, and after 11 days, neural crest cells were passaged onto fibronectin-coated 6-well plates (∼1m cells/well) using accutase and fed with neural crest maintenance media (NMM). NMM: 1:1 ratio of DMEM-F12 and neurobasal, 0.5x Gem21 NeuroPlex Supplement with Vitamin A (Gemini, 400-160), 0.5x N2 NeuroPlex Supplement (Gemini, 400-163), 1x antibiotic/antimycotic, 0.5x Glutamax, 20ng/ml bFGF, 20ng/ml bFGF EGF and 1mg/ml BSA (Gemini). After 2-3 days, neural crest cells were plated at ∼1m cells/well of a 6-well plate and the following day cells were fed with neural crest long-term media. Long term media: neural crest maintenance media + 50pg/ml BMP2 (PeproTech, 120-02) + 3uM CHIR-99021 (Selleck Chemicals, S2924) (BCh media). After transition to BCh media, CNCCs at subsequent passages were plated at ∼800,000 cells/well of a 6-well plate. CNCCs were then passaged twice to passage 4 where depletion experiments were performed, or cells were further differentiated to chondrocytes. For depletion experiments, dTAG^V^-1 (Tocris, 6914) at a range of concentrations was added to BCh media, with an equivalent amount of DMSO as vehicle control.

To differentiate CNCCs to chondrocytes, passage 3 CNCCs were passaged to passage 4, seeded at ∼250,000 CNCCs per well of a 12 well plate, and grown for 3 days in BCh media. Then, CNCCs were transitioned to chondrocyte media without TGFb3 (ChM), with or without dTAG^V^-1. ChM: DMEM-HG, 5% FBS, 1x ITS premix, 1mM sodium pyruvate, 50 μg/mL ascorbic acid, 0.1 μM dexamethasone and 1x antibiotic/antimycotic. The following day, cells were fed with chondrocyte media with TGFb3 (ChMT), with or without dTAG^V^-1. ChMT: ChM + 10 ng/mL TGFb3. Cells were fed every subsequent 3 days with ChMT. Cells were harvested at day 10 and/or day 21 of the differentiation.

### Sulfated glycosaminoglycan (sGAG) quantification

Total sGAG levels per well of chondrocytes independently differentiated from CNCCs for 21 days, representing mature cartilage formation, were quantified using the Blyscan glycosaminoglycan assay (Biocolor). Briefly, collagen in the extracellular matrix (ECM) was digested by washing cells with PBS and then adding 1 mL of Papain digestion buffer per well of a 12-well plate. Cells were incubated at 65C for 3 hours with gentle agitation every 30 min, then 0.5 mL additional digestion buffer was added and lysate was moved to Eppendorf tubes and incubated at 65C overnight. Quantification of sGAG content from ∼10 uL of the lysates was performed as per manufacturer instructions, and the volume of each lysate was measured separately and used to infer the total sGAG content of the entire well.

### Flow cytometry

CNCCs were harvested for flow cytometry using accutase and quenching with FACS buffer (5% FBS in PBS). Chondrocytes were harvested as described previously with the following modifications. Chondrocytes were incubated in digestion medium for ∼1hr with gentle agitation every 15 min. Digestion medium: DMEM-KO, 1mg/mL Pronase (Roche, 11459643001), 1mg/mL Collagenase B (Roche, 11088815001), 4U/mL Hyalauronidase (Sigma, H3506-500mg). Digested cells were then washed twice in PBS. Flow cytometry was used to measure mNeonGreen fluorescence after excluding doublets and debris based on forward and side scatter (Beckman Coulter Cytoflex). Fluorescence values were summarized per biological replicate using geometric means. The relative SOX9 level as % of the *SOX9*-tagged, unperturbed (treated with DMSO) sample was calculated by first subtracting the geometric mean fluorescence of the untagged (WT) sample from both the unperturbed and dTAG^V^-1-treated sample, then dividing the dTAG^V^-1-treated sample fluorescence by the unperturbed sample fluorescence.

### Protein harvesting and western blotting

Cells were washed with PBS and scraped into RIPA buffer, incubated on ice for 10 min, sonicated to disrupt pelleted DNA using Bioruptor Plus (Diagenode). RIPA buffer: 50 mM Tris, 150 mM NaCl, 1% NP-40, 0.1% Na-deoxycholate, 0.1% SDS in H20 with 1x protease inhibitor cocktail (Sigma-Aldrich 4693132001). Sonicated lysates were incubated one ice for 10 min, and centrifuged at 16,000 RCF for 10 min at 4C to pellet debris. Supernatants were normalized to the same protein content using Pierce BCA Protein Assay kit (ThermoFisher, 23225), mixed with 4x SDS sample loading buffer (Invitrogen NP0007) and 0.1M DTT, and boiled for 7 minutes. Samples were separated on Tris-glycine PAGE gels in 1x tris-glycine buffer with 0.1% SDS, transferred in 1x tris-glycine buffer with 20% methanol, blocked in 5% milk + 1% BSA in PBST, immunoblotted with either SOX9 antibody (1:1000, Sigma-Aldrich AB5535) or beta-Actin antibody (1:20,000, Abcam ab49900) overnight at 4C, probed with the appropriate secondary, developed using Pierce ECL Western Blotting Substrate (ThermoFisher, 32106), and imaged using an Amersham ImageQuant 800 system (Cytiva).

### RNA isolation and preparation of RNA-seq libraries

Total RNA was extracted from CNCCs using Trizol reagent (Invitrogen) followed by Quick-RNA Miniprep kit (Zymo) with on-column DNase I digestion. Unstranded mRNA libraries were prepared with the NEBNext Ultra II RNA Library Prep Kit for Illumina (NEB #E7770S/L).

### Metabolic RNA labeling and preparation of SLAM-seq libraries

Metabolic RNA labeling and SLAM-seq was performed as previously described ^42^, with the following modifications. 4-Thiouridine (4SU) was incorporated into nascent transcripts by incubating CNCCs with BCh media containing 100 uM 4SU, as well as DMSO or 500 nM dTAG^V^-1 depending on experimental condition, for 2 hours. Plates were covered in foil and handling was done in a hood with no light where possible. For 3 and 24 hour depletion experiments, labeling was started at 1 and 22 hours after dTAG^V^-1 addition, respectively.

Total RNA was extracted using Trizol reagent, phenol-chloroform extraction was performed, and the aqueous phased was used as input to the Quick-RNA Miniprep kit. During RNA extraction with Quick-RNA Miniprep kit, 0.1 mM DTT was added to the the RNA wash and RNA pre-wash buffers, but the on-column DNase I step was skipped. RNA was eluted in H20 with 1 mM DTT, quantified with Qubit RNA Broad Range assay (ThermoFisher, Q10211), and ≥ 2 ug total RNA was used as input to the alkylation reaction. Alkylation was performed in dark tubes after which light exposure was allowed, and after quenching RNA was purified and subjected to on-column DNase I digestion using the RNA Clean & Concentrator-5 kit (Zymo, R1013).

500 ng alkylated RNA was used as input to QuantSeq 3’ mRNA-Seq Library Prep Kit FWD with unique dual index (UDI) add-on (Lexogen, 113.96), with 15 cycles of PCR amplification. Library size distributions were confirmed by separation on a PAGE gel and staining with SYBRGold and pooled based on quantifications from Qubit dsDNA High Sensitivity Kit (ThermoFisher , Q32854). Pooled libraries were sequenced using Novaseq 6000 platform (2x 150bp).

### ATAC-seq harvesting and library preparation

ATAC-seq was performed as described previously^59^. Briefly, CNCCs were incubated with BCh media containing 200U/mL DNaseI (Worthington, LS002007) for 30 min AND harvested using accutase. Viable cells were counted using Countess Automated Cell Counter (Invitrogen), 50,000 viable cells were pelleted at 500 RCF for 5 min at 4°C and resuspended in ATAC-resuspension buffer containing 0.1% NP40, 0.1% Tween20, and 0.01% Digitonin and incubated on ice for 3 minutes. Following wash-out with cold ATAC-Resuspension Buffer (RSB, 10 mM Tris-HCl pH 7.4, 10mM NaCl, 3mM MgCl2 in sterile water) containing 0.1% Tween20, cells were pelleted and resuspended in 50 μL transposition mix (25 μL 2x TD buffer, 2.5 μL transposase (100nM final), 16.5 μL PBS, 0.5 μL 1% digitonin, 0.5 μL 10% Tween20, 5 μL H_2_O) and incubated for 30 minutes at 37°C with shaking. The reaction was purified using the Zymo DNA Clean & Concentrator kit and PCR-amplified with NEBNext master mix and primers as defined in Corces et al. Libraries were purified by two rounds of double-sided size selection with AMPure XP beads (Beckman Coulter, A63881), with the initial round of 0.5x sample volume of beads followed by a second round with 1.3x initial volume of beads. Library size distributions were confirmed by separation on a PAGE gel and staining with SYBRGold and pooled based on quantifications from Qubit dsDNA High Sensitivity Kit. Pooled libraries were sequenced using Novaseq 6000 platform (2x 150bp).

### Chromatin immunoprecipitation (ChIP) and library preparation

One fully confluent 10 cm plate of cells was cross-linked per ChIP experiment in 10 mL PBS with 1% methanol-free formaldehyde for 10 min and quenched with a final concentration of 0.125M glycine for 5 min with nutation. Cross-linked cells were scraped into tubes with 0.001% Triton X in PBS, washed with PBS without Triton, pelleted by centrifugation, flash-frozen in liquid nitrogen and stored at −80°C. Samples were defrosted on ice and resuspended in 5mL LB1 (50 mM HEPES-KOH pH 7.5, 140 mM NaCl, 1 mM EDTA, 10% glycerol, 0.5% NP-40, 0.25% Triton X-100, with 1X cOmplete Protease Inhibitor Cocktail and PMSF) and rotated vertically for 10 min at 4°C. Samples were centrifuged for 5 min at 1350 x g at 4°C, and resuspended in 5mL LB2 (10 mM Tris, 200 mM NaCl, 1 mM EDTA, 0.5 mM EGTA, with 1X cOmplete Protease Inhibitor Cocktail and optionally 1mM PMSF) and rotated vertically for 10 min at 4°C. Samples were centrifuged for 5 min at 1350 x g at 4°C, resuspended in 300 uL LB3 per sonicated sample, and incubated for 10 min on ice. Samples were sonicated in 1.5 mL Bioruptor Plus TPX microtubes (Diagenode, c30010010-50) on Bioruptor Plus for 10 cycles of 30sec ON/30sec OFF. Every 5 cycles, samples were lightly vortex and briefly centrifuged. Samples were diluted in additional LB3 to 1 mL, pelleted at 16,000 RCF for 10 min, and the supernatant was removed. Triton X-100 was added to 1%.

To check DNA size distribution and quantity, a 10 uL aliquot of sonicated chromatin from each sample was diluted to 100 uL in Elution Buffer (50 mM Tris, 10 mM EDTA, 1% SDS) with 0.0125 M NaCl and 0.2 mg/mL RNase A and incubated at 65C for 1 hour, followed by addition of Proteinase K to 0.2 mg/mL and an additional 1 hour of 65C incubation. DNA was purified using Zymo DNA Clean & Concentrator Kit with ChIP DNA Binding Buffer (Zymo, D5201-1-50) and size distribution and quantity was assessed by separation on a 1% agarose gel and Qubit HS DNA kit, respectively. Qubit measurements were used to normalize samples to the same DNA concentration.

Following normalization, the chromatin was divided for input (2%) and ChIP samples. A minimum of 25 ug DNA was used for histone ChIPs, and 50 ug for V5 ChIPs. 5 μg anti-H3K27ac (Active Motif, 39133) antibody or 10 μg anti-V5 (Abcam, ab9116) antibody was added per ChIP sample, and incubated overnight at 4°C. Protein G Dynabeads (ThermoFisher) were first blocked with Block solution (0.5% BSA (w/v) in 1X PBS) and then added to cleared chromatin to bind antibody-bound chromatin for a 4-6 hour incubation. Chromatin-bound Dynabeads were washed at least 6 times with chilled RIPA wash buffer (50 mM HEPES-KOH pH 7.5, 500 mM LiCl, 1 mM EDTA, 1% NP-40, 0.7% Na-Deoxycholate), followed by a wash with chilled TE + 50 mM NaCl. Chromatin was eluted for 30 min in Elution Buffer (50 mM Tris, 10 mM EDTA, 1% SDS) at 65°C with frequent vortexing. The ChIP and input samples were then incubated at 65°C overnight to reverse cross-links (12-16 hours). Samples were diluted and sequentially digested with RNase A (0.2 mg/mL) for 2 hours at 37°C followed by Proteinase K (0.2 mg/mL) for 2 hours at 55°C for 2-4 hours to digest protein. ChIP and input samples were purified by Zymo DNA Clean & Concentrator Kit with ChIP DNA Binding Buffer.

For library preparation, samples were quantified by Qubit dsDNA HS assay kit, and 10-50ng of ChIP DNA was used for library preparation with end repair, A-tailing, and adaptor ligation (NEB). Following USER enzyme treatment, libraries were cleaned up with one round of single-side AMPure XP bead clean-up, then amplified to add indices using NEBNext Ultra II Q5 Master Mix and NEBNext Multiplex Oligos for Illumina kit (NEB, E7335S) with 4-10 cycles (as determined by input amounts from NEB protocol). ChIP libraries were purified by two rounds of double-sided AMPure XP bead clean-up (0.5x then 0.4x initial sample volume of beads added) to remove large fragments and deplete adaptors. Library concentration and quality within ChIP or input groups was assessed by Qubit dsDNA HS assay kit and separation on a PAGE gel, and used to pool within ChIP or input groups. KAPA qPCR was used to pool across ChIP or input groups. Pooled libraries were sequenced using Novaseq 6000 platform (2x 150bp).

### Sequencing data pre-processing

#### ATAC-seq and ChIP-seq

For both ATAC-seq and ChIP-seq, Nextera (ATAC) or Truseq (ChIP) adapter sequences and low-quality bases (-Q 10) were trimmed from sequencing reads using skewer v0.2.2 and aligned to the human genome (hg38) using bowtie2 v2.4.1 with the following settings: --very-sensitive, --X 2000. Read mate pair information was corrected with samtools v1.10 fixmate, PCR duplicates were removed using samtools markdup, and mitochondrial reads and low mapping quality reads (-q 20) were removed using samtools view. bigWig files for visualization were generated using deeptools v3.5.0 bamCoverage with the following settings: -bs 10 -- normalizeUsing RPGC --samFlagInclude 64 --samFlagExclude 8 --extendReads.

For ATAC-seq, a custom approach based on a previous publication ^17^ was used to define regions that showed reproducible peaks of accessibility across samples. Shifted bed sites were obtained from mapped and filtered ATAC bam files, and bed files for each sample were used to call peak summits using MACS2 v2.2.7.1 callpeak with the following settings: --nomodel --keep-dup all --extsize 200 --shift 100 --SPMR. Then, within each differentiation/line replicate, summits within 75bp were merged, taking the average location across summits as the location of the merged summit. Then, across each differentiation/line, summits within 150 bp were merged, again taking the average location. Only those merged summits with at least one constituent summit from three or more differentiation/line instances were carried forward. These summits were extended 250bp in either direction, and finally all such regions were merged such that there were no overlapping regions, resulting in 151,457 reproducible peak regions.

#### RNA-seq

TruSeq adapter sequences and low-quality bases were trimmed from sequencing reads using skewer, and transcript levels were quantified using salmon v1.4.0 quant with the following settings: --gcBias --seqBias -l A. Salmon abundance files were summarized to the gene level and imported into R with the tximport package v1.20.0 with countsFromAbundance = “lengthScaledTPM”.

#### SLAM-seq

Lexogen adapter sequences and low-quality bases were trimmed from sequencing reads (read 1 only) using skewer, followed by trimming of poly(A) sequences. Trimmed reads were used as input to slamdunk v0.4.3^60^, with the following individual step parameters modified from default: map, -n 100 -5 0; count, -l 150.

### Quantification and statistical analysis

#### Sequence motif matching

Transcription factor (TF) sequence motif position weight matrices (PWMs) for the indicated TFs were obtained from HOCOMOCO core motifs: SOX9, SOX9_HUMAN.H11MO.0.B; TFAP2A, AP2A_HUMAN.H11MO.0.A; NR2F1, COT2_HUMAN.H11MO.0.A. The Coordinator motif corresponding to TWIST1 was obtained from a previous publication ^38^. The SOX9 palindrome motif was constructed by inverting the single HOCOMOCO PWM at various spacings from 0-10 bp. All motifs were matched to the human genome using fimo v5.1.1 with a p-value threshold of 1e-4.

#### Differential expression/accessibility testing

Differential expression or accessibility between pairs of SOX9 concentrations (ATAC/RNA) or timepoints of full SOX9 depletion (ATAC, SLAM, H3K27ac ChIP) was performed using DESeq2 v1.32.0, with CNCC differentiation batch as a covariate and raw counts as input. For SLAM one additional surrogate variable, discovered using sva 3.4.0, was also used as a covariate. For ATAC and H3K27ac ChIP, counts over all 151,457 reproducible peak regions were used, for RNA only protein-coding genes with at least 1 transcript per million (TPM) in at least 6 samples were used, and for SLAM-seq only protein-coding genes with at least 1 count per million (CPM) in at least 3 samples were used. The independentFiltering option in DESeq2 was set to FALSE, except for H3K27ac differential analyses.

#### Modeling of SOX9 dose-response curves (ATAC/RNA)

All RE/gene CPM values were first TMM-normalized using the edgeR package. For each SOX9-dependent RE/gene, defined by 5% FDR comparing depleted vs fully depleted SOX9, CPM values across all *SOX9*- tagged samples (i.e. from all six SOX9 concentrations) were corrected for differentiation batch effect by linear regression using the lm() function. Differentiation-corrected CPM values were scaled by dividing by the maximum absolute value across samples. Outliers, defined as z-score greater than 3, were removed. The data were then fit to either a linear model as a function of SOX9 dosage (defined by flow cytometry), or to Hill equation using the drm() function in the drc package. For most genes/REs, a two-parameter Hill equation (i.e. with minimum and maximum fixed as the mean CPM at full or no depletion, respectively) was sufficient.

However, for a small subset of genes/REs, a three-parameter Hill equation with fixed minimum but free maximum was a better fit (decrease in AIC > 2); for these genes/REs, the three-parameter Hill was used. To calculate the ‘buffering index’ at a given SOX9 dosage such as 50% (see Extended Data Fig. 3), the change in the fitted Hill equation curve going from 100% to 50% SOX9 dosage was divided by the total SOX9-dependent change (i.e. going from 100% to 0%), multiplied by 100, then subtracted from 100. A value of 0 of this statistic indicates no buffering (i.e. the entirety of SOX9-dependent change has occurred by 50% SOX9 dosage) while a value of 100 indicates complete buffering (i.e. no change until < 50% SOX9 dosage).

#### Bootstrapping for ED50/Hill exponent confidence interval estimation

Point estimates for ED50 and Hill exponent from fitted Hill equations vary nonrandomly with both the relative quality of the fitted Hill equation (with fitted parameters for REs/genes fit better by a linear model having more uncertainty) as well as the overall magnitude of ED50/Hill exponents (higher magnitudes having greater uncertainty). We noticed instability in the ED50/Hill standard errors obtained from parametric least-squares fitting in the drc package; we therefore implemented a bootstrap procedure to quantify uncertainty in ED50/Hill estimates at either the individual RE/gene level or when comparing groups of REs/genes in their ED50 or Hill exponent values. For each RE/gene, a set of 200 bootstrapped datasets was generated by sampling the number of replicates (generally 7) with replacement from each of the six conditions. Note that while the number of potential bootstraps from a single condition is relatively small (7!), performing this sampling independently in each of the six conditions generates a very large number of unique datasets (7!^6^). Hill equations were fit to each bootstrapped dataset and ED50/Hill exponents were extracted.

For uncertainty estimates for individual genes, the 200 bootstrap replicates were summarized to determine 95% confidence intervals. When comparing groups of genes, rather than first summarizing bootstraps within genes, the relative group statistic (typically median) was computed across all genes for each of 200 bootstrap replicates separately; the resulting 200 group statistics were then used to construct 95% confidence intervals.

#### Prediction of SOX9-dependent RNA changes from ATAC changes

An extension of the Activity-by-Contact (ABC) model ^44^ was used to predict gene expression fold-changes at each SOX9 concentration (relative to undepleted) from ATAC-seq fold changes at nearby REs from the same comparisons. Briefly, the (ABC) model defines the contribution, or ABC score, of a given RE within 5Mb of a gene TSS as:

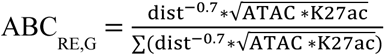

ABC scores for all RE-gene pairs (within 5Mb) were calculated using this formula. In this case a linear distance-power law function was used as a proxy for ‘Contact,’ as it has been shown to perform similar to Hi-C ^44^. A gene’s own promoter (defined as an RE within 1 kb of the consensus TSS) was excluded for the purposes of gene-level predictions, since promoter accessibility is often reflective of gene transcriptional changes. For ‘Activity’ calculations, ATAC-seq and H3K27ac counts from unperturbed (*SOX9*-tagged, DMSO-treated) CNCCs were used.

A gene’s predicted relative level at a certain SOX9 concentration was calculated as the sum of the ABC scores of all REs within 5Mb. Since H3K27ac ChIP-seq was only available from unperturbed or fully depleted *SOX9*- tagged CNCCs, RE ABC scores at lower SOX9 concentrations were calculated by multiplying the unperturbed ABC score by the DESeq-estimated fold-change for that RE when comparing unperturbed CNCCs to the given SOX9 concentration. While this assumes an identical decrease in H3K27ac at every SOX9 concentration, fold-changes in RE ATAC and H3K27ac signals were observed to be highly correlated upon full SOX9 depletion. Effectively, this approach predicts the fold-change in gene expression as a weighted sum of fold-changes in all REs within 5Mb, where the weights are the RE ABC scores from the unperturbed setting:

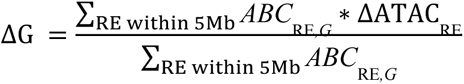

### Pierre Robin Sequence (PRS) endophenotype definition and GWAS

#### Sample

The control sample of healthy individuals comprised three-dimensional facial scans of 8,246 unrelated individuals of European ancestry (60.3% female; median age = 18.0 years, IQR = 9.0 years) originating from the US and the UK. A detailed description of the sample is provided in a previous publication ^30^. The sample of Pierre Robin Sequence comprised 13 participants (9 female; median age = 12.01 years, IQR = 5.17 years). Images were excluded if participants were laughing, crying or otherwise emoting or judged to be of poor quality or if the non-rigid registration failed. Participants with missing covariate information (*e.g.* age, sex) were additionally removed.

#### Genotyping

Imputed genotypes were available for all individuals of the European control sample, as described in detail in a previous publication ^30^. After quality control, 7,417,619 SNPs were used for analysis. SNPs on the X chromosome were coded 0/2 for hemizygous males, to match with the 0/1/2 coding for females.

#### Phenotyping

##### Correction for asymmetry and covariates

Facial images were processed in Meshmonk to obtain a standard facial representation, characterized by 7160 homologous quasi-landmarks including midline and bilaterally-paired quasi landmarks^61^. Each configuration was made symmetrical following the Klingenberg protocol^62^: for each configuration a reflected copy was made by reversing the sign of the x coordinate of each quasi landmark. Bilaterally-paired quasi-landmarks were relabeled left to right and right to left in the reflected copy. The reflected and relabeled copy was then aligned to the original by least-squares Procrustes superimposition. The average of the two copies was taken as the symmetrical version of the configuration.

The US and UK samples were adjusted for covariates sex, age and age-squared as follows. All symmetrized quasi-landmark configurations were aligned by generalized Procrustes analysis. The average configuration was recorded. A partial least-squares regression of the configurations onto the covariates was performed. The average configuration was added to the residuals to produce the corrected configurations of the US and UK samples. The regression coefficients were retained to adjust the PRS sample for the same covariates using the same regression model. Specifically, each symmetrized landmark configuration of the PRS sample was aligned to the recorded average configuration. The predicted configuration for their sex, age and age-squared was calculated from the recorded regression coefficients and was subtracted from their symmetrized and aligned configuration. The coordinates of the average configuration were then added back on to produce the corrected version of the PRS participant.

##### PRS-driven phenotyping

Facial shape was partitioned into 63 global-to-local segments by hierarchical spectral clustering as previously described (Claes et al., 2018). For each subset of quasi-landmarks belonging to each of the 63 facial segments, a PRS-driven univariate trait was defined as follows. First the symmetrized and adjusted quasi-landmark configurations of the US and UK samples were co-aligned by generalized Procrustes analysis, and this for each segment separately. The dimensionality was reduced by principal component analysis with the optimal number of principal components to retain determined by parallel analysis. Projections on each principal component were normalized to have unit variance by dividing each projection by the standard deviation of all projections. These standard deviations were retained. The symmetrized and adjusted landmark configurations of the PRS sample were then aligned to the average and projected into the space of the principal components and normalized by the recorded standard deviations. Finally, per facial segment, a PRS-driven facial trait was defined as the vector or direction passing through the global average and average PRS facial shape.

Each participant in the US and UK samples was ‘scored’ on the PRS-driven facial traits by computing the cosine of the angle between: 1) the vector from the average of the PC projections of the US and UK samples to the PC projections of the participant and 2) the vector from the average of the US and UK projections to the average of the PRS projections. These scores were computed by leave-one-out such that each participant was excluded from training the vectors on which they were scored.

##### Significance testing

To test the significance of the PRS-driven trait in each facial module, the sample of PRS were compared to a matched control sample of equal size drawn from the US and UK samples. The matched control sample was selected randomly as follows, separately for each facial module. In random order, each participant in the PRS sample was matched to the participant from the combined US and UK samples of the same sex that was closest in age. This participant was then removed from the possible matches so that each US/UK participant could only be matched to one PRS participant. The covariate-adjusted and symmetrized quasi-landmarks were co-aligned by generalized Procrustes analysis and regressed onto group membership (0=US/UK; 1=PRS) using partial least-squares regression. A p-value was generated by a permutation test on R-squared with 10,000 permutations. In 30 out of 63 facial segments, a significant difference (*p* ≤ 0.05) in facial shape was observed between the two groups (PRS vs healthy controls).

#### GWAS

The scores on the 30 PRS-driven univariate traits, for which a significant difference was observed, were combined into a single phenotype matrix ([N x M] with N = 8246 controls and M = 30 facial segments). This matrix was tested for genotype-phenotype associations in a multivariate meta-analysis framework using canonical correlation analysis (CCA), similar to the work of White et al. (2021). However, instead of performing a separate GWAS per facial segment, information across multiple segments is now combined into a single multivariate GWAS. Because CCA does not accommodate adjustments for covariates, we removed the effect of relevant covariates (sex, age, age-squared, height, weight, facial size, four genomic ancestry axes, camera system), on both the independent (SNP) and the dependent (facial shape) variables using partial least-squares regression prior to GWAS.

The US and UK subsamples served both as identification and replication sets in a two-stage design, after which the p-values were meta-analyzed using Stouffer’s method^63, 64^. Per SNP, the lowest p-value was selected (meta_US_ vs meta_UK_) and compared against the genome-wide Bonferroni threshold (5 x 10^-8^). We observed 1767 SNPs at the level of genome-wide significance, which were clumped into 22 independent loci as follows. Starting from the lead SNP (lowest p-value), SNPs within 10kb or within 1Mb but with r^2^>0.01 were clumped into the same locus represented by the lead SNP. Next, considering the lead SNPs only, signals within 10Mb and an r^2^>0.01 were merged. Third, any locus with a singleton lead SNP was removed.

---

**Extended Data Figure 1.**
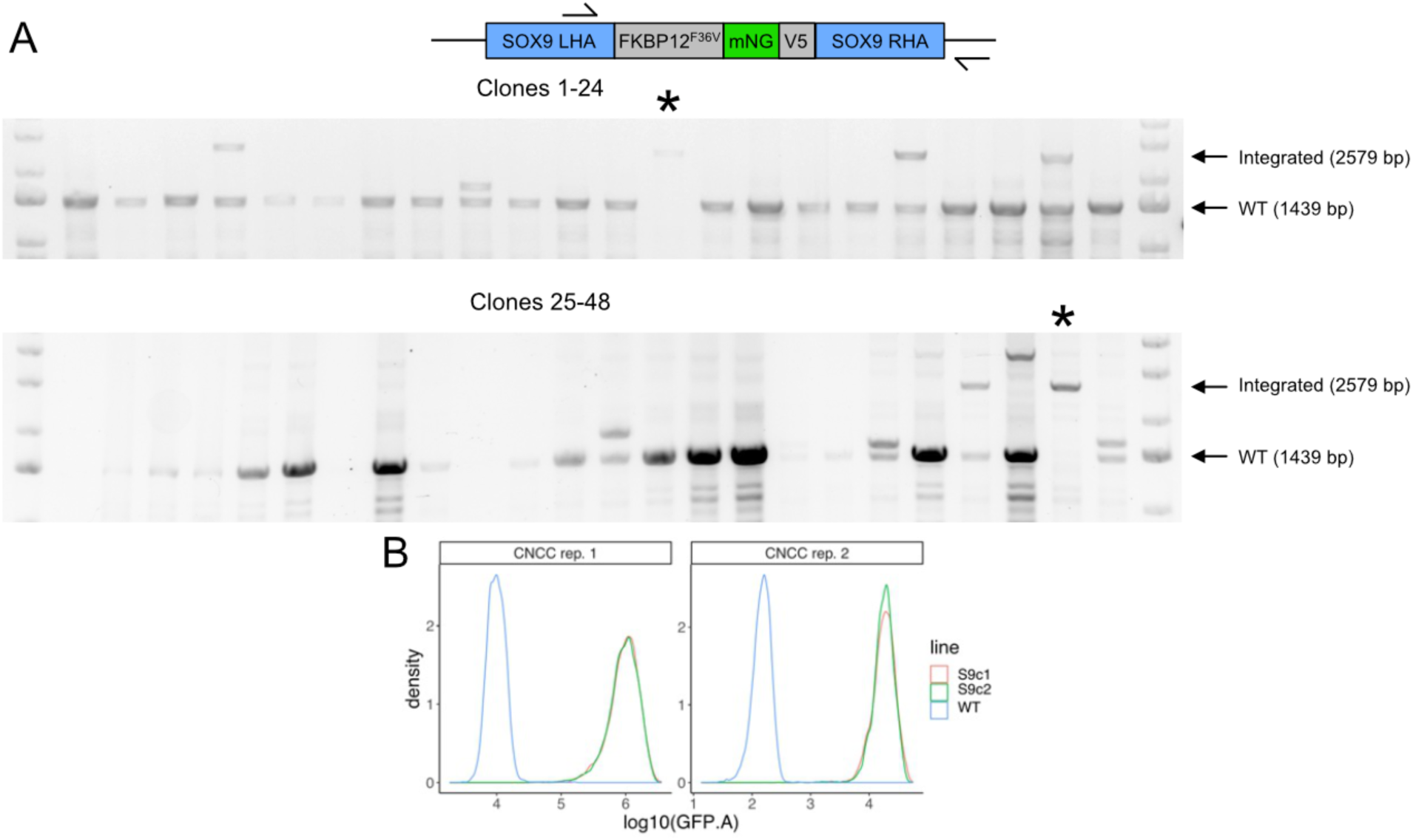
(A) Genome editing of WT hESCs to derive SOX9-tagged hESCs. Top, schematic depicting primer (arrow) locations for clonal genotyping of the *SOX9* locus. SOX9LHA-FKBP12^FV36^-mNG-V5-SOX9RHA is the full homology-directed repair template provided by AAV6, so the right primer is located outside the homology arm. Bottom, agarose gel images of PCR using depicted primers on 48 analyzed hESC clones nucleofected with *SOX9* sgRNA-Cas9 RNP and transduced with tag-containing AAV6. * clones with bi-allelic knock-in. (B) Single-cell distributions of mNeonGreen fluorescence between two SOX9-tagged clones from two CNCC replicates.

**Extended Data Figure 2.**
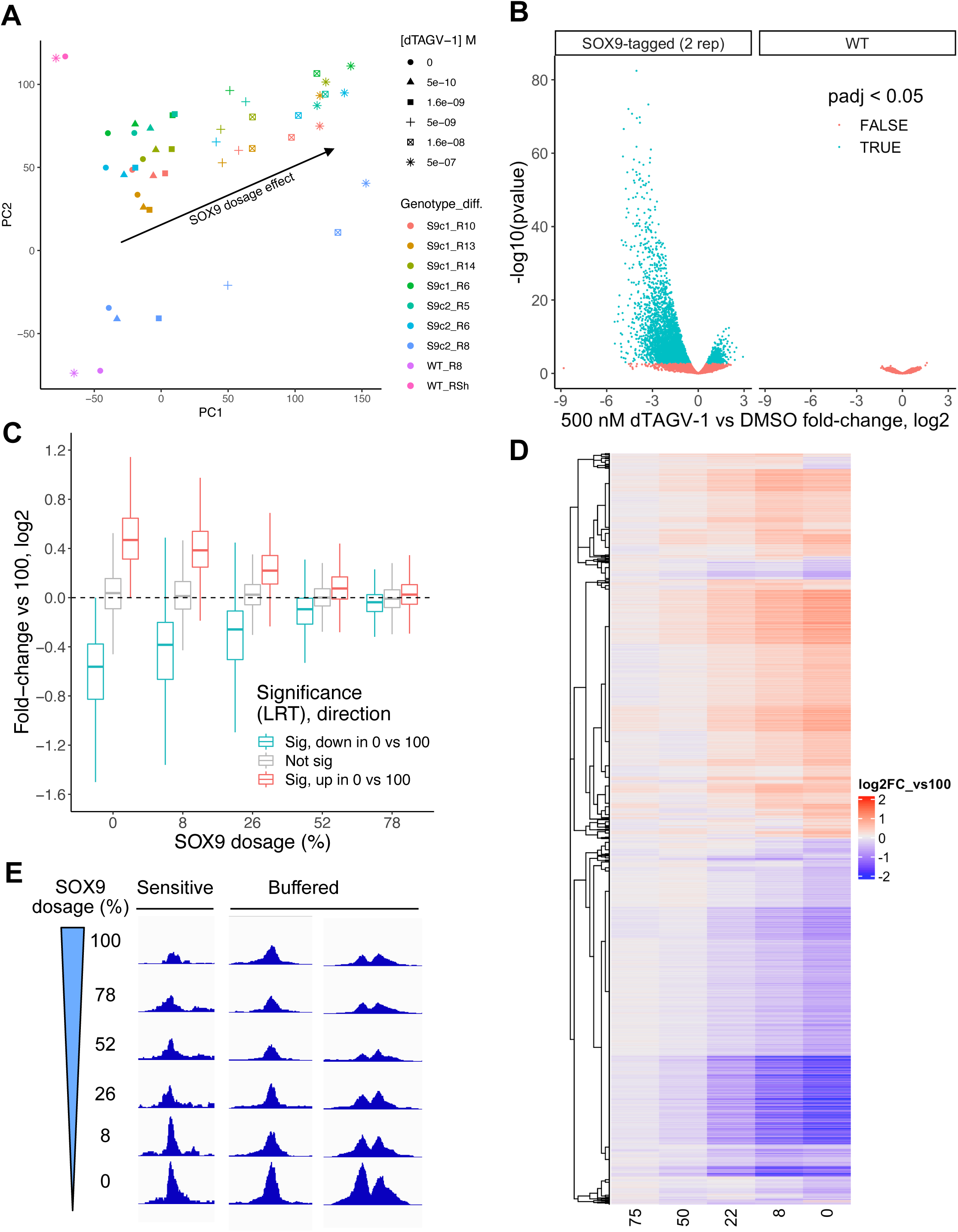
(A) Principal component analysis of ATAC-seq counts per million (CPM) of all 151,457 REs across all CNCC samples. Shapes indicate the dTAG^V^-1 concentration treated for 48h. Colors indicate the combination of hESC line from which CNCCs were derived and differentiation batch (S9c1/2 = *SOX9*-tagged clone1/2). Arrow indicates the SOX9 dosage effect. (B) Volcano plot of 500 nM dTAG^V^-1 treatment on two SOX9-tagged (left) or two WT (right) CNCC differentiation replicates for all 151,457 REs. (C) Distributions of fold-changes versus full (100) SOX9 dosage for all genes for which SOX9 dosage explains a significant (5% FDR, red, blue) or nonsignificant (grey) amount of variance (likelihood ratio test, LRT), stratified by the direction of change in full SOX9 depletion. (D) Same fold-change values as in (C) for a random subset (10,000) of significant REs, plotted as a heatmap and clustered by row based on Kendall distance. (E) Example ATAC-seq browser tracks of individual RE accessibility at different SOX9 dosages (y-axis), averaged across all replicates at each dosage.

**Extended Data Figure 3.**
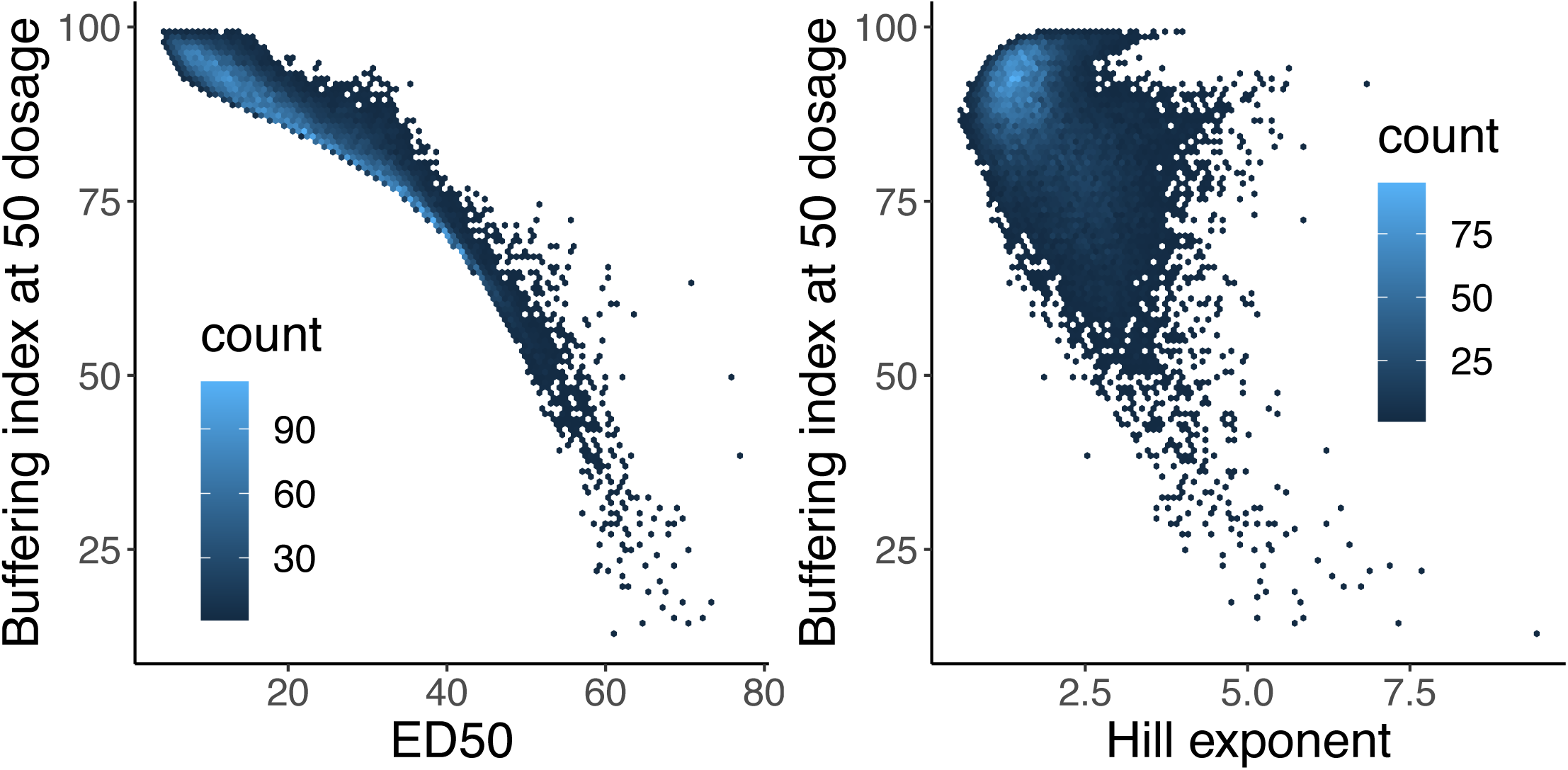
For all SOX9-dependent REs with good Hill equation fits (p < 0.05 for both ED50 and Hill exponent), correlation between either ED50 (left, Spearman ρ -0.961) or Hill exponent (right, Spearman ρ -0.457) and buffering index calculated at 50% SOX9 dosage. See Methods for details on calculation of buffering index – 0 means no buffering (effect of 100 to 50% SOX9 dosage on RE accessibility is 50% of effect of 100 to 0% SOX9 dosage), 100 means full buffering (no effect of 100 to 50% SOX9 dosage on RE accessibility, but significant effect of 100 to 0% SOX9 dosage).

**Extended Data Figure 4.**
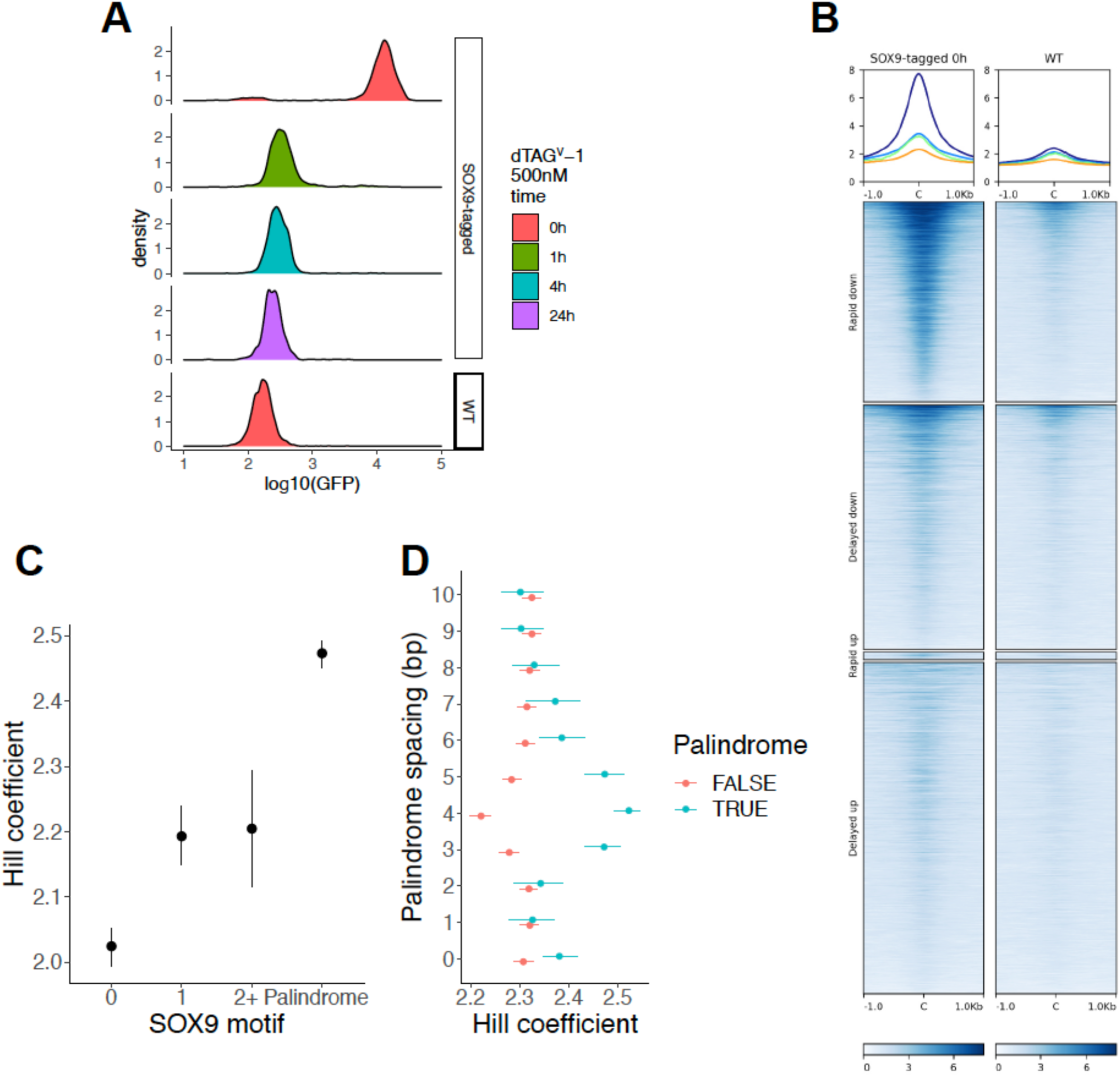
(A) mNeonGreen fluorescence intensity in SOX9-tagged or WT CNCCs with the times of treatment by dTAG^V^-1. (B) V5 ChIP-seq signal from CNCCs with V5-tagged SOX9 present (“*SOX9*-tagged 0h”) or absent (“WT”) plotted over sets of SOX9-dependent REs as defined in Fig. 3A. (C) Hill exponent of rapid down REs stratified by SOX9 motif type, with motif position weight matrices as in Fig. 3E. (D) For the SOX9 palindrome motif at with a 0-10 bp spacing between the inverted repeats (y-axis), rapid down REs were stratified on the basis of that motif match. Points and error bars represent median and 95% confidence intervals as computed by bootstrap (see Methods).

**Extended Data Figure 5.**
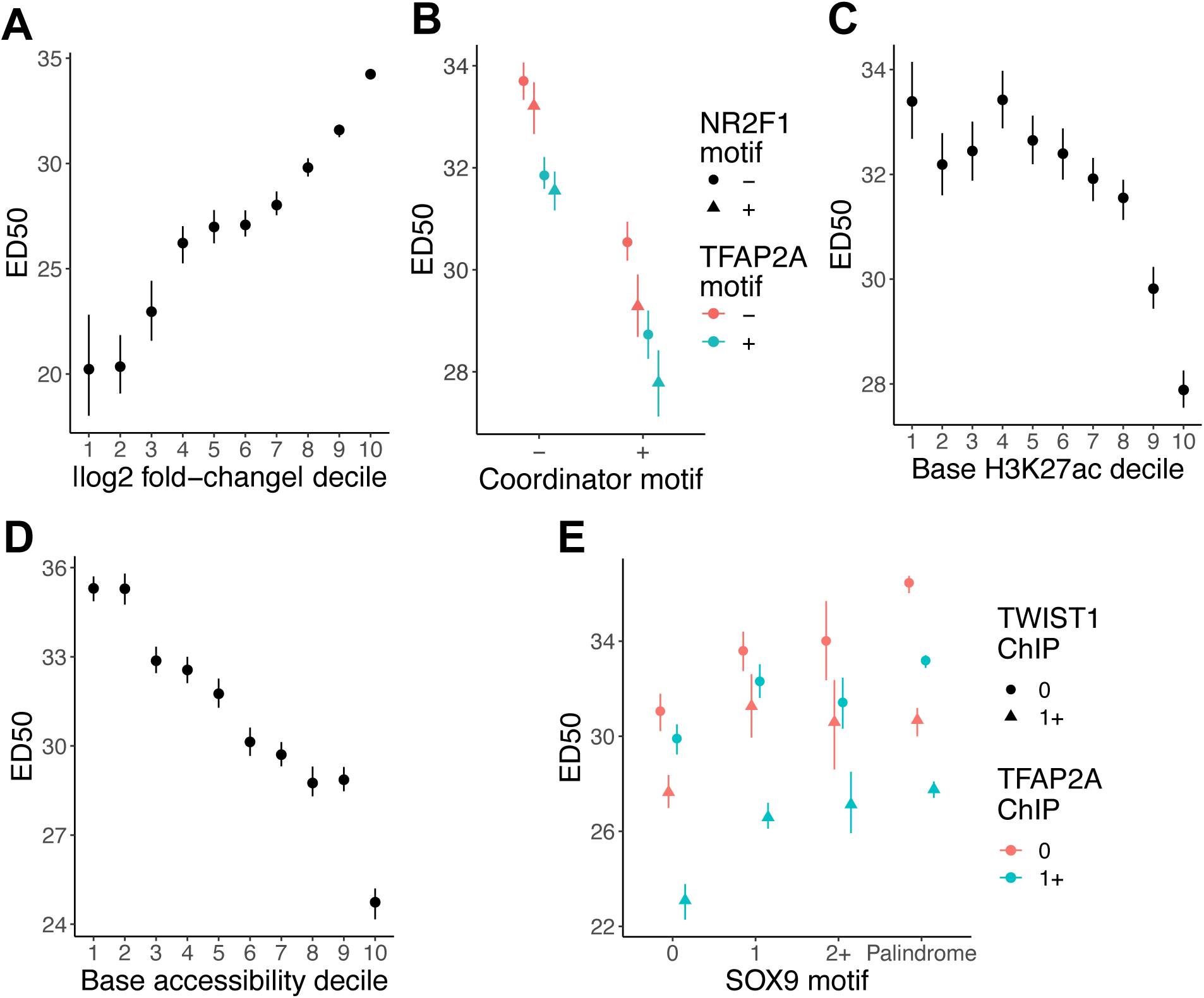
**Additional features affecting sensitivity of the RE response to SOX9 dosage changes.** ED50 of rapid down SOX9-dependent REs, stratified by (A) magnitude of change in response to full SOX9 depletion, (B) presence of Coordinator (TWIST1), NR2F1, or TFAP2A sequence motif matches, baseline levels of (C) H3K27ac or (D) chromatin accessibility, or (E) the combination of SOX9 motif type and TWIST1/TFAP2A binding by ChIP-seq. For decile plots, higher deciles mean higher values. Points and error bars represent median and 95% confidence intervals as computed by bootstrap (see Methods).

**Extended Data Figure 6.**
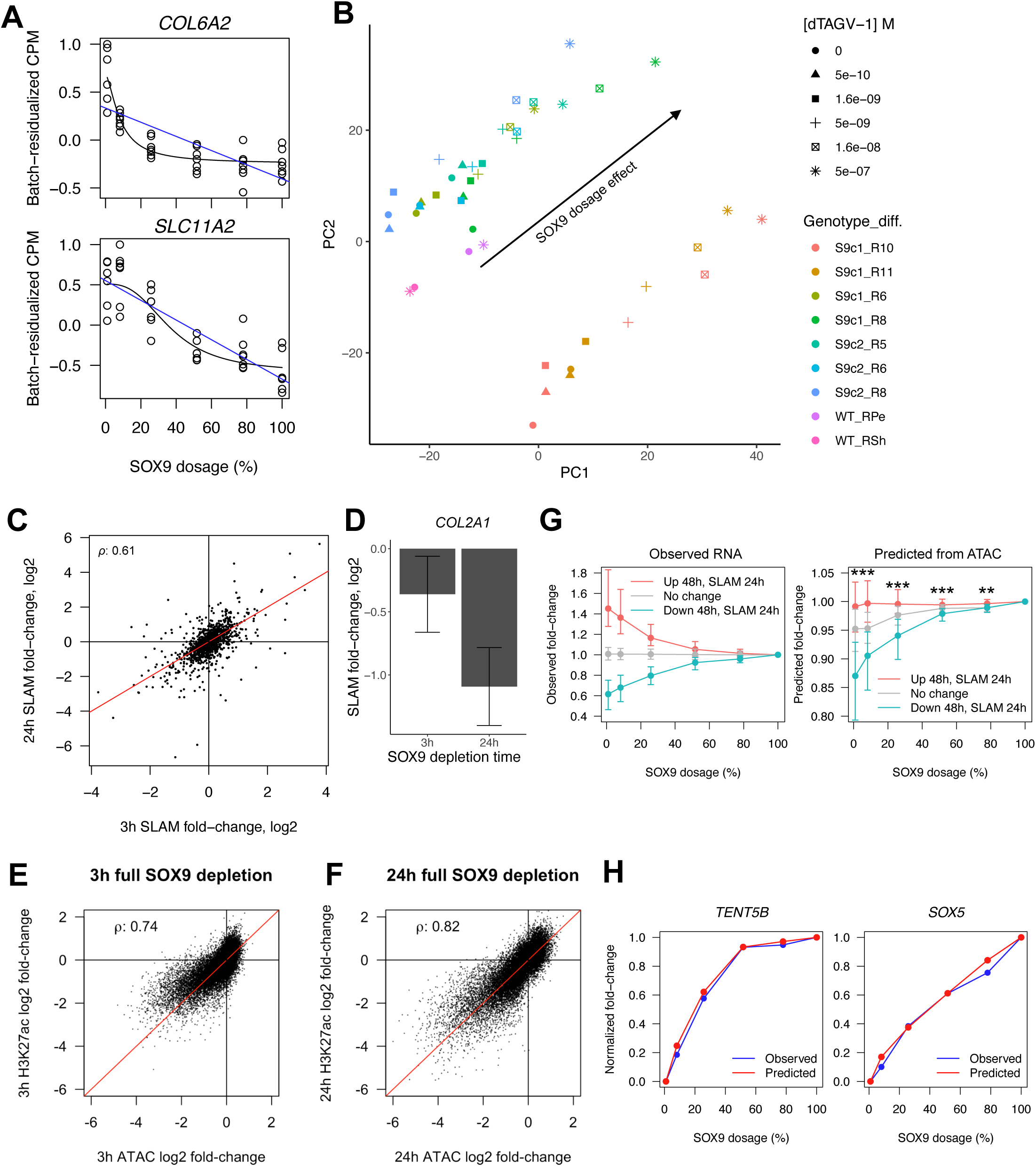
(A) Examples of genes upregulated in response to SOX9 depletion with buffered (top) or switch-like (bottom) responses. (B) Principal component analysis of RNA-seq counts per million (CPM) across all SOX9-tagged and WT CNCC samples. Shapes indicate the dTAG^V^-1 concentration treated for 48h. Colors indicate the combination of hESC line from which CNCCs were derived and differentiation batch (S9c1/2 = SOX9-tagged clone1/2). Arrow indicates the SOX9 dosage effect. (C) Scatterplot of effects of full SOX9 depletion for 3h (x-axis) versus 24h (y-axis) on nascent transcription, as assayed by SLAM-seq, for all SOX9-dependent genes (i.e. responding to full SOX9 depletion for 48h in RNA-seq). Y=x line in red. (D) Effects of full SOX9 depletion for 3h (left) or 24h (right) on transcription of *COL2A1*, a known direct target of SOX9. Barplot represents point estimate of log_2_ fold-change from DESeq2, error bars represent 95% confidence interval. (E,F) For all REs responding to full depletion of SOX9 at 48h, the effects of 3h (E) or 24h (F) full SOX9 depletion on chromatin accessibility (ATAC-seq, x-axis) or H3K27ac levels (ChIP-seq, y-axis) is plotted. (G) Distributions of observed (left) or predicted (right) fold-changes vs full SOX9 dosage at each concentration, stratified based on direction of transcriptional response to full SOX9 depletion (colors). N of groups by color: red, 184; grey, 10,339; blue, 197. Points and error bars represent median and 25^th^ and 75^th^ percentiles of distribution. (H) Examples of predictions for a buffered (left) or sensitive (right) gene. ** p < 1e-6, *** p < 2.2e-16, Wilcoxon rank-sum test.

**Extended Data Figure 7.**
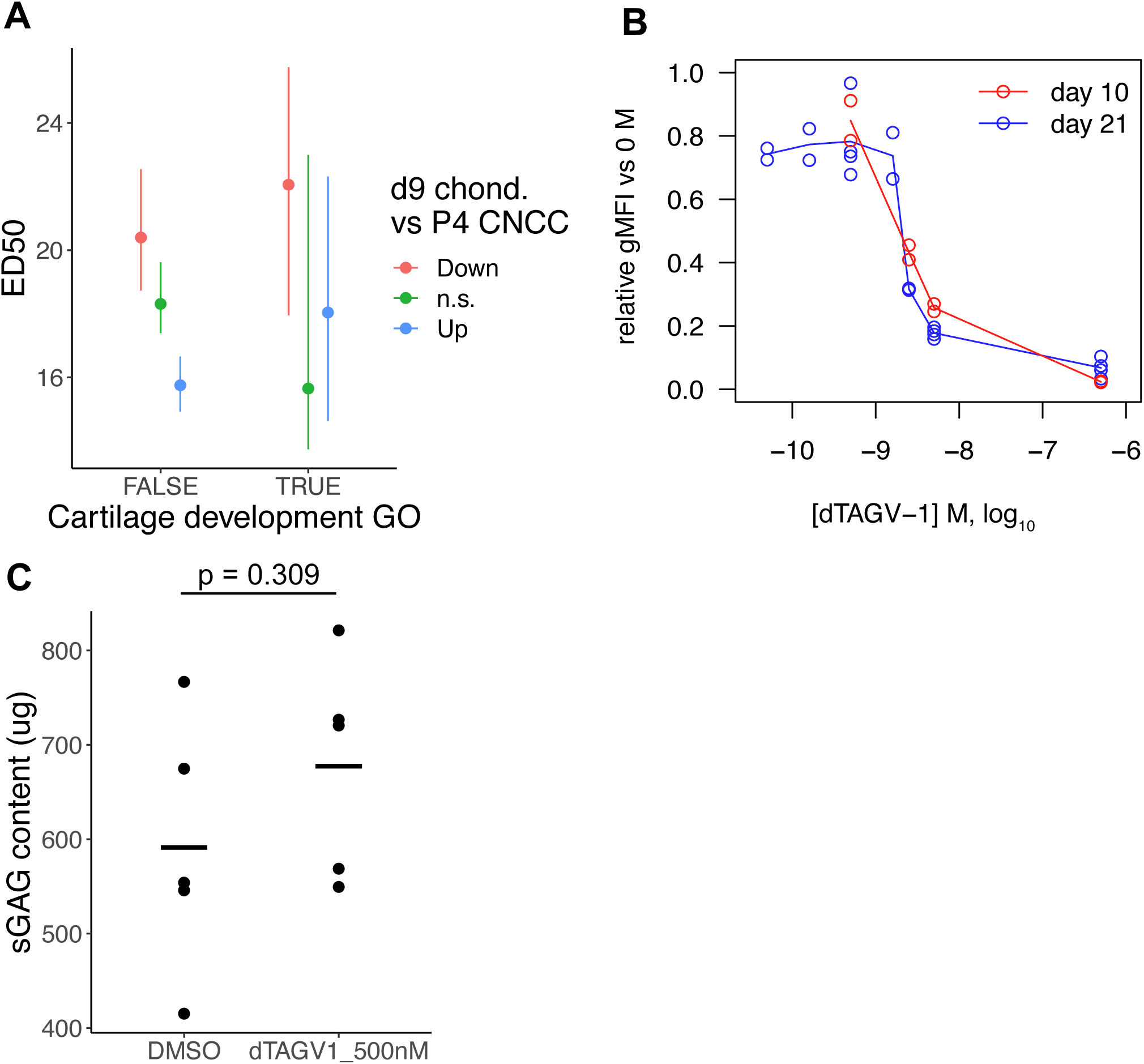
(A) ED50 of SOX9-upregulated genes stratified by presence in the “Cartilage development” Gene Ontology (GO) category (x-axis), and expression change in chondrocytes compared to CNCCs (color, data from Long et al 2020). N of groups from left to right: 94, 217, 204, 3, 9, 17. Points and error bars in (A,E) represent median and 95% confidence intervals as computed by bootstrap (see Methods). (B) Fluorescence intensity at day 10 (red) or 21 (blue) of chondrogenesis in SOX9-tagged chondrocytes as a function of dTAG^V^-1 concentration. gMFI, geometric mean. (C) Sulfated glycosaminoglycan (sGAG, representative of mature cartilage) at day 21 of chondrogenesis in WT CNCCs treated with DMSO or 500 nM dTAG^V^-1. Bars represent mean, p-value from two-sided T-test.

**Extended Data Figure 8.**
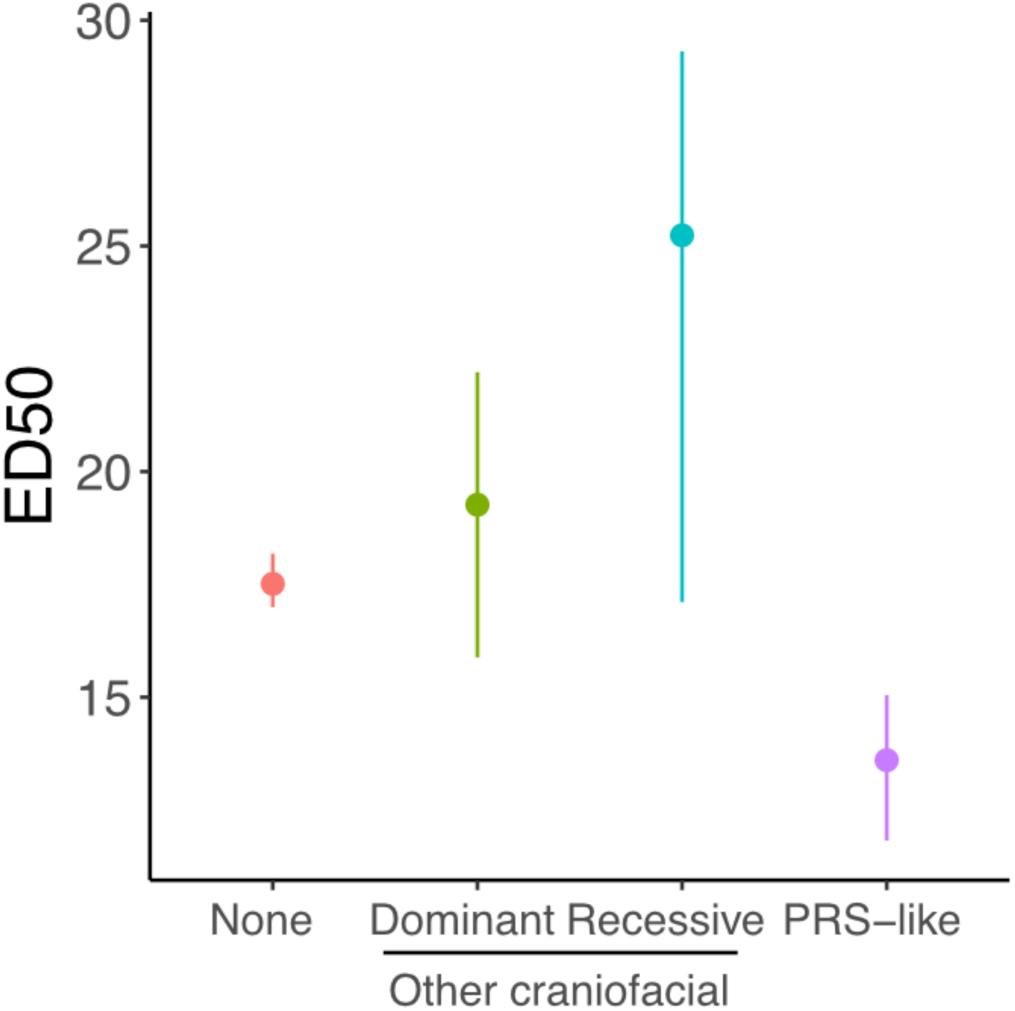
ED50 by craniofacial disorder association for genes upregulated upon SOX9 depletion. Gene-craniofacial disorder associations determined as in Figure 6A. N of groups from left to right: 508, 20, 9, 5. Points and error bars represent median and 95% confidence intervals as computed by bootstrap (see Methods).

**Extended Data Figure 9.**
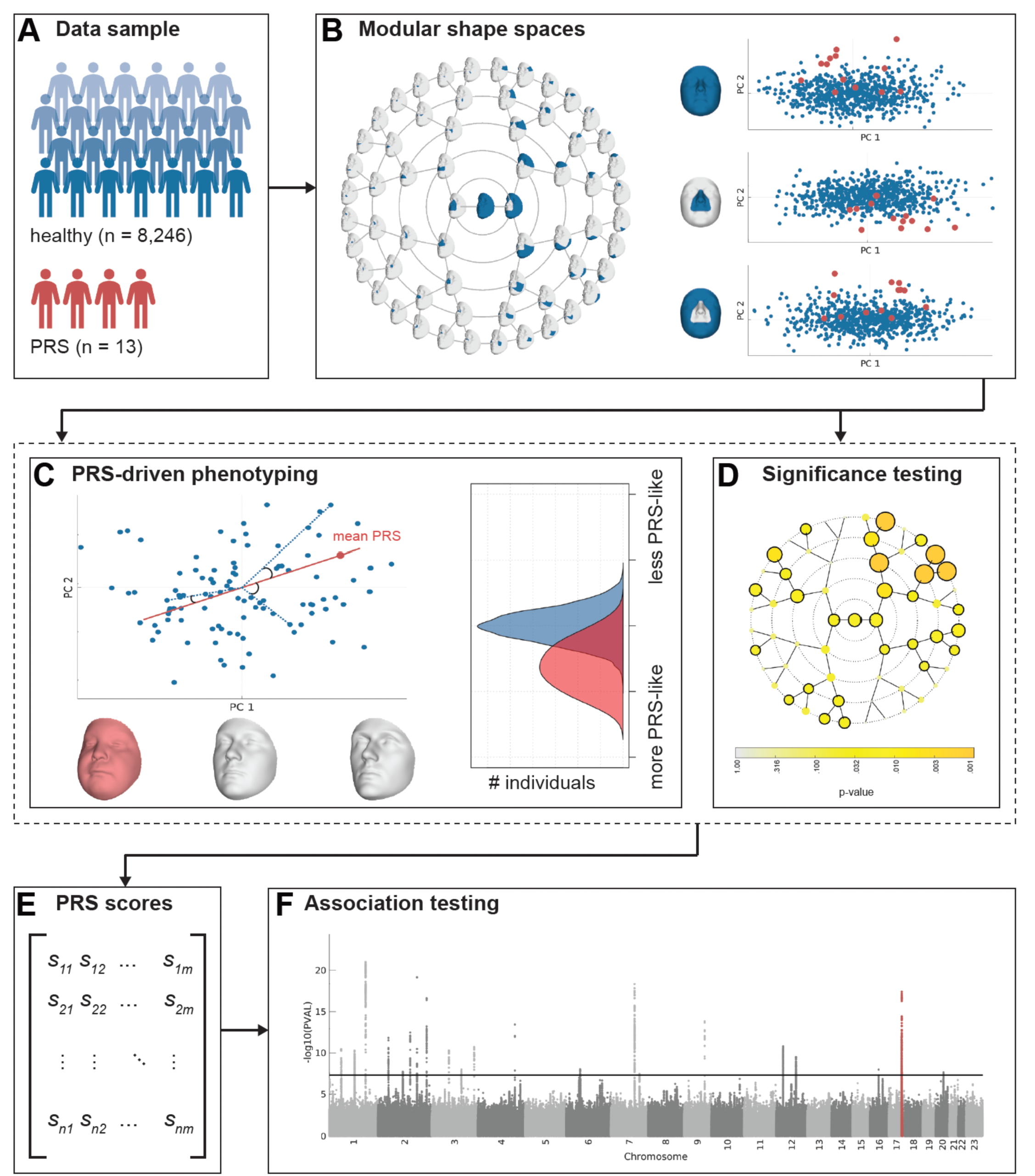
**Schematic of endophenotype definition approach and GWAS in healthy individuals.** (A) The study sample consisted of 8,246 healthy, unrelated European-ancestry individuals and 13 patients with Pierre Robin Sequence (PRS). (B) Global-to-local segmentation of 3D facial shape obtained using hierarchical spectral clustering of the European cohort. For each of the facial segments (n = 63) a shape space is established based on the larger European cohort (blue dots) using PCA, describing the main axes of variation in the data. The PRS facial shapes (red dots) are then aligned and projected onto each segment-derived PCA space. (C) Per facial segment, a PRS-driven univariate trait is defined as the vector passing through the global European average facial shape (center) and the PRS average (red dot). Each trait or direction (red line) represents a complex shape transformation that codes for PRS-characteristic facial features, as displayed by the three facial morphs (right = typical PRS face; middle = average face; left = opposite or anti-face). In a leave-one-out approach, each individual was scored on the PRS-driven facial traits by computing the cosine of the angle between the vector going from the global European average to each participant (blue dotted lines), and the vector from the global European average to the average PRS projection (red line). Scores range from 0 to 2, with scores close to 0 indicating the presence of facial features similar to those typically observed in PRS, whereas scores close to 2 correspond to features opposite to PRS. (D) To test the significance of the PRS-driven trait in each facial segment, the sample of PRS were compared to a matched control sample of equal size drawn from the larger European cohort using partial least squares regression and a p-value was generated by a 10,000-fold permutation test. In 30 out of 63 facial segments a significant difference (p<=0.05, black encircled segments) was observed between the PRS sample and healthy controls. (E) The scores on each of the 30 significant traits were combined into a single phenotype matrix ([8246 x 30]) (F) and subsequently tested for genotype-phenotype associations in a multivariate GWAS meta-analysis approach using canonical correlation analyses. Association statistics per SNP are displayed in the Manhattan plot, with the region surrounding *SOX9* highlighted in red.

**Extended Data Figure 10.**
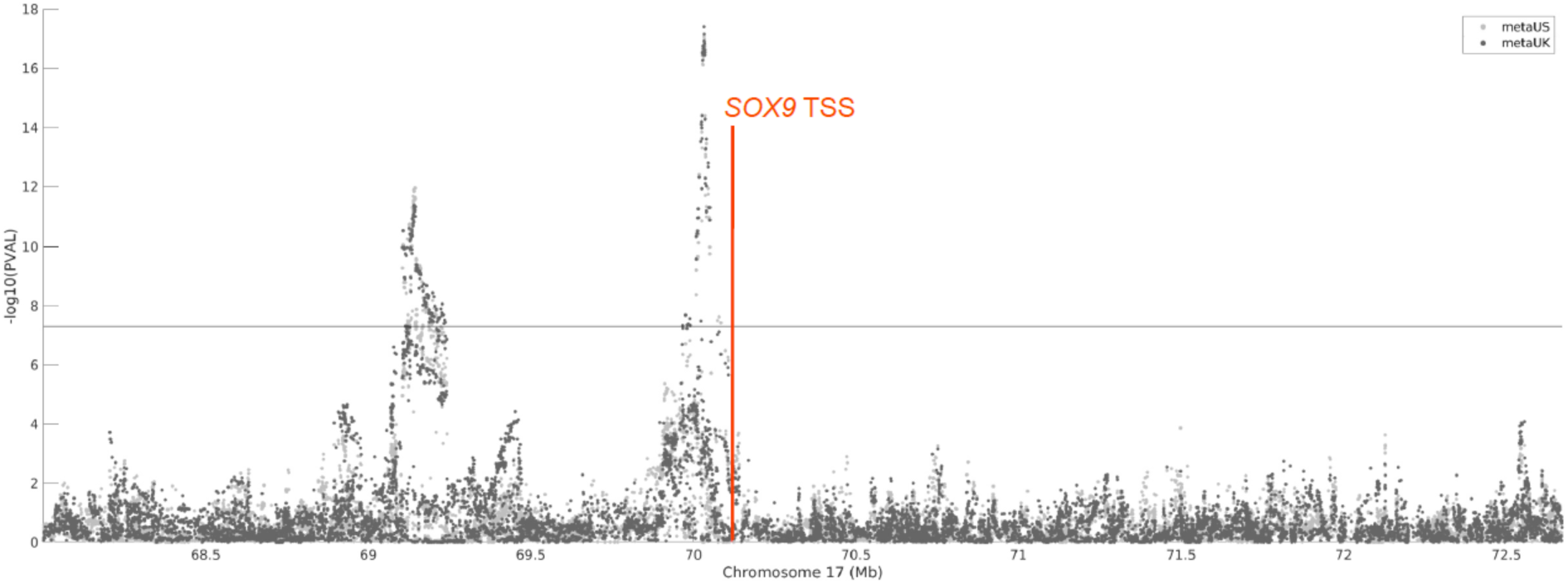
**Association of variants near *SOX9* with PRS endophenotypic variation in healthy individuals.** Plot of -log10(p-value) (y-axis) versus SNP position (x-axis) in two different cohorts (color). Location of the *SOX9* transcription start site (TSS) is indicated in red. Horizontal line represent genome-wide significance.

## SUPPLEMENTARY TABLES

**Supplementary Table 1. Additional information for all SOX9-dependent REs.**

**Supplementary Table 2. Additional information for all SOX9-dependent genes.**

**Supplementary Table 3. All genome-wide significant lead SNPs from PRS endophenotype GWAS.**

